# High energy charge, antioxidant capacity, and amino acid contents are conserved features of T cell metabolism

**DOI:** 10.1101/2025.06.09.658709

**Authors:** Samantha O’Keeffe, Harin Sim, Elle McCue, Junyoung O. Park

**Affiliations:** Department of Chemical and Biomolecular Engineering, University of California, Los Angeles, Los Angeles, CA 90095, USA; Department of Bioengineering, University of California, Los Angeles, Los Angeles, CA 90095, USA; Department of Mechanical and Aerospace Engineering, University of California, Los Angeles, Los Angeles, CA 90095, USA

## Abstract

T cell diversity and differentiation define individual adaptive immunity. Metabolism fuels adaptive immunity by driving proliferation, differentiation, and activation; however, how individual and lineage differences shape T cell metabolism remains unknown. Here we develop an ensemble method for absolute metabolome quantitation and quantify 84 metabolites in human primary T cells. Liquid chromatography-mass spectrometry of metabolites co-extracted from T cells and ^13^C-labeled reference cells reveals absolute concentrations *en masse*. Across subtypes and individuals, T cell metabolomes resemble one another. T cells possess thermodynamically forward-driven glycolysis, high adenylate energy charge, large amino acid pools, and favorable redox ratios for anabolism and antioxidant defense. Across metabolism, metabolite concentrations exceed their associated Michaelis constants and inhibitor constants two thirds and half of the time, respectively. The conserved features of T cell metabolomes underlie a design principle: metabolite levels prime T cells for increased energy demand, oxidative stress, and rapid regulation that accompany immune response.

---

Individual variations in the immune system give rise to different levels of protection against diseases and have implications for developing immunotherapy^1-3^. In the context of T cells in adaptive immunity, individual variations arise from genetic diversity^4^, T cell receptor (TCR) diversity^5^, and differentiation into subtypes^6^. Since metabolism converts nutrients into usable energy and biochemical building blocks necessary for T cell differentiation, proliferation, and function^7-10^, a fundamental question arises: how do T cell diversity and differentiation manifest in metabolism?

T cells are maintained in a quiescent state through tightly regulated metabolic processes^11^ to promote survival and proliferation without activation^12^. T cell metabolism is reprogrammed in response to antigen stimulation^13^ and sometimes mere expression of surface proteins such as chimeric antigen receptors (CARs) ^14^. Among broad changes, energy generation shifts from oxidative phosphorylation to aerobic glycolysis, amino acid uptake increases, and overflow metabolism commences^14,15^. Such metabolic reprogramming brings about an increased supply of ATP, amino acids, and nucleotides for effective immune response on one hand^15^, and increased reactive oxygen species (ROS)^16,17^ and altered redox homeostasis that contribute to exhaustion and autoimmune disease pathogenesis on the other hand^18,19^.

These prior findings paint relative changes in T cell metabolism as immune biomarkers and therapeutic targets^10,11^; however, a baseline metabolic profile of T cells is necessary to establish a solid foundation for mechanistic understanding and rational tuning of the adaptive immune system.

Absolute metabolite concentrations dictate cell physiology (e.g., signaling and osmolarity) and metabolic rates through Michaelis-Menten kinetics (**Eqn. 1**) and thermodynamics (**Eqn. 2**). For a reaction of the form S⇌P, its rate is described by the reversible Michaelis-Menten equation:

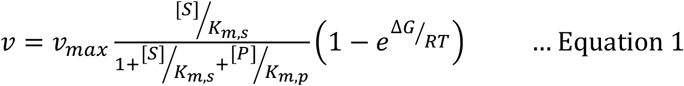

v_m*ax*_ is the maximum reaction rate, [S] and [P] are the concentrations of substrate S and product P, K_m,s_ and K_m,p_ are the Michaelis constants for S and P, R is the gas constant in J mol^−1^ K^−1^, T is the temperature in Kelvins, and ΔG the Gibbs free energy of the reaction. ΔG is modified from its value at biochemical standard conditions (ΔG°’) to account for the effect of metabolite concentrations:

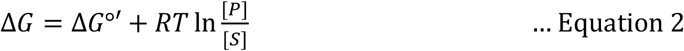

Cells regulate metabolite levels to efficiently utilize metabolic networks and avoid toxic buildup^20^. In turn, metabolites shape metabolic activity by acting as substrates, products, and effectors^21-24^. The value of absolute metabolite concentrations warrants comprehensive metabolome quantitation in T cells.

Liquid chromatography-mass spectrometry (LC-MS) has been a workhorse in metabolomics^25^. Challenges in quantifying absolute metabolite concentrations in human T cells are twofold: *i*) biologically, primary T cell availability is limited for each individual; and *ii*) analytically, current detection tools have varying response factors (signal-to-quantity ratios) for different metabolites. To address the latter analytical challenge, cellular metabolites are spiked with known concentrations of internal standards. Stable isotopes such as ^13^C are used to distinguish between intracellular metabolites and internal standards (**Fig. 1a**).

**Figure 1.**
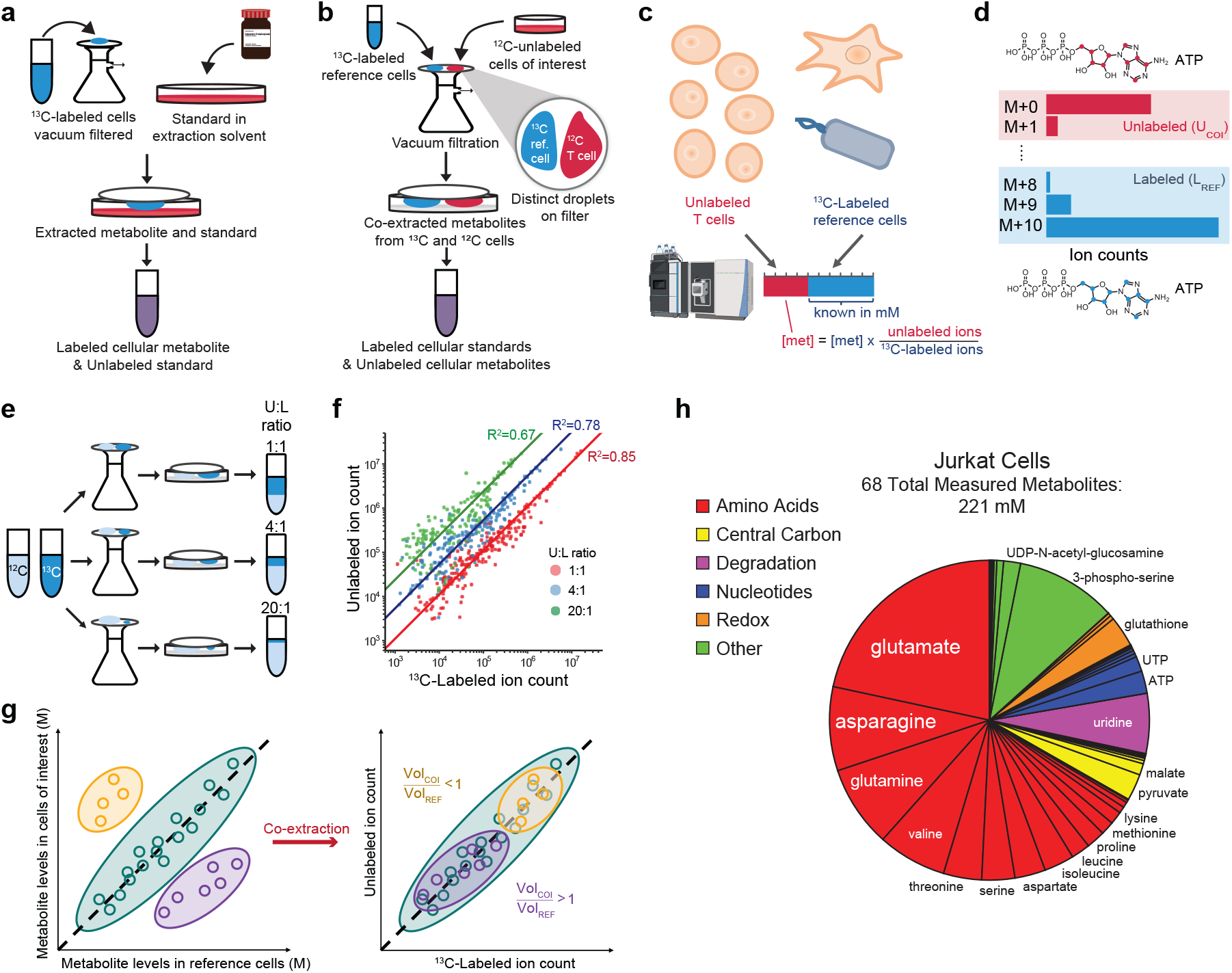
Metabolite co-extraction enables ensemble quantitation using ^13^C-labeled metabolome as internal standards. (**a**) Current absolute metabolite quantitation relies on the use of an authenticated standard dissolved in metabolite extraction solvent. Cells of interest are grown in ^13^C-labeled media and extracted into the solvent containing the known concentration of the unlabeled standard. (**b**) In the present work, co-extraction of metabolites from two cell types, one of which was ^13^C-labeled, enabled absolute quantitation of multiple metabolites in parallel. Care was taken to not put the two cell types in contact during co-extraction. (**c**) The resulting LC-MS peaks contained a mixture of unlabeled metabolites from T cells and ^13^C-labeled metabolites from reference cells. Using the ratios of unlabeled to labeled metabolite peaks and the known concentrations of metabolites in reference cells (blue), absolute metabolite concentrations in T cells were obtained. (**d**) For each metabolite that has n carbons, multiple mass isotopomers with monoisotopic masses from M+0 (unlabeled) to M+n (fully ^13^C-labeled) were observed. With natural ^13^C abundance, isotope impurity, and ^12^CO_2_ incorporation metabolic pathways in mind, each mass isotopomer was attributed to either the cell of interest (COI) or the reference cell (REF). (**e**) This ensemble method was tested by co-extracting unlabeled and ^13^C-labeled *E. coli* at the ratios of 1:1, 4:1 and 20:1. (**f**) The accuracy of the method was validated by comparing the measured ratios of the unlabeled (U) to labeled (L) ion counts to the expected U:L ratios (n=3 biological replicates for each ratio) for individual metabolites. (**g**) Cell density and extraction volumes were tuned to achieve U:L ratios near 1:1 to minimize measurement errors. (**h**) Upon co-extractions with ^13^C-labeled *E. coli* and iBMK cells, Jurkat T cell metabolites were quantified using the ensemble method.

Known concentrations of internal standards and LC-MS measurement of labeled-to-unlabeled isotope ratios for individual metabolites reveal their absolute concentrations^26,27^. The present isotope ratio-based approach, while effective for a handful of metabolites or for immortalized cell lines, does not overcome the dual challenges of comprehensive absolute metabolome quantitation in human primary T cells.

To address these challenges, we developed an ensemble metabolite quantitation method for absolute quantitation of metabolite concentrations in human primary T cells. The ensemble method leverages the known metabolite concentrations in model systems including *E. coli* and immortalized mammalian cells^24^ as internal reference. The reference cells were grown on uniformly ^13^C-labeled glucose, [U-^13^C_6_]glucose, and co-extracted with T cells whose metabolite concentrations had been unknown (**Fig. 1b**). The known concentrations of metabolites in the reference cell and LC-MS measurement of labeled-to-unlabeled isotope ratios revealed absolute metabolite concentrations *en masse* (**Fig. 1c**).

Using the ensemble method, we quantified the absolute concentrations of 84 metabolites totaling 131-154 mM and 221 mM in human primary T cells and Jurkat cells, respectively. To our surprise, absolute metabolite concentrations displayed a strong resemblance across subtypes and individuals. Amino acids were the largest class of metabolites in T cells with glutamate and aspartate being the most abundant metabolite in Jurkat cells and primary T cells, respectively. The energy status of T cells was healthy with a high adenylate energy charge of 0.95 and favorable redox ratios NADH/NAD^+^≈0.06 and NADPH/NADP^+^≈19. Thermodynamic analysis revealed that glycolysis and glutathione reductase were moderately to strongly forward driven with Gibbs free energy changes (ΔG) of –67 kJ/mol and –30 kJ/mol, respectively. Kinetic analysis revealed that T cell metabolomes facilitated efficient enzyme utilization and regulation. These findings pointed to a metabolic design principle: conserved metabolite levels prime T cells for energy generation, antioxidant activity, and efficient regulation. Thus, our ensemble quantitation established a reference human T cell metabolome and the healthy baseline state of T cell metabolism.

## Results

### Co-extraction enables absolute metabolome quantitation *en masse*

Absolute metabolite concentrations offer quantitative insights into metabolic regulation, energetics, and enzyme efficiency, but a demand for large numbers of cells and internal standards forbid their measurement in human primary cells . To overcome these challenges, we innovated absolute metabolome quantitation in primary human T cells that uses isotopically labeled cellular metabolites as internal standards by co-extracting metabolites from the cells of interest (COI) and ^13^C-labeled reference cells (REF). For each metabolite, ^12^C-dominant peaks and ^13^C-dominant ions were divided into unlabeled and labeled groups to account for ^12^C impurities and natural ^13^C abundance (**Fig. 1d**). During co-extraction, two cell types were kept separate to prevent unwanted interactions and perturbations until they were quenched (**Fig. 1b**). By using reference cells with known concentrations^24^, the co-extraction method obviated the need for individual standards and parallelized absolute quantitation of multiple metabolites in a single sample.

To validate our approach, we performed co-extraction with known ratios of two cultures of *E. coli* grown on unlabeled or ^13^C-labeled glucose in otherwise identical conditions (**Fig. 1e**). LC-MS analysis revealed the ratios of unlabeled to ^13^C-labeled metabolites (U:L ratio) as expected from the ratios of extracted culture volumes (**Fig. 1f**). At all ratios tested (U:L ratio equal to 1:1, 4:1, and 20:1), the measurements globally stayed near the expected ratio with R^2^=0.85, R^2^=0.78, and R^2^=0.67, respectively. However, the distributions of measured U:L ratios became broader as they deviated further from one due to the lower limit of detection and ion suppression in the mass spectrometer (**Extended Data Fig. 1**). Since one-to-one ratios of unlabeled to ^13^C-labeled metabolites minimized variance and error, we tuned the U:L ratios to be near one by co-extracting different amounts of the COI and the REF (**Fig. 1g**). Collectively, ^13^C-labeling, judicious co-extraction, and LC-MS enabled parallel quantification of metabolites from limited cells. In Jurkat T cells, we quantified 68 metabolite concentrations, totaling 221 mM (**Fig. 1h**).

### Conservation of T cell metabolome across individuals

We conducted absolute metabolome quantitation in human primary T cells from individual healthy donors. To best capture *in vivo* T cell metabolism in two major T cell subtypes, we isolated CD4^+^ and CD8^+^ T cells by negative selection from peripheral blood mononuclear cells (PBMCs) because ligand binding on cell surface may affect cellular metabolism^14^. To allow T cells to reach homeostasis, we incubated them for two hours in RPMI 1640 medium supplemented with 10% dialyzed fetal bovine serum. Within seven to nine hours of blood collection, we extracted CD4^+^ and CD8^+^ T cell metabolites using the ensemble method (**Fig. 2a**). The rapid extraction and measurement of T cell metabolome minimized the alteration of T cell proteome to reveal *in vivo*-like T cell metabolism before it adapted to *in vitro* conditions.

**Figure 2.**
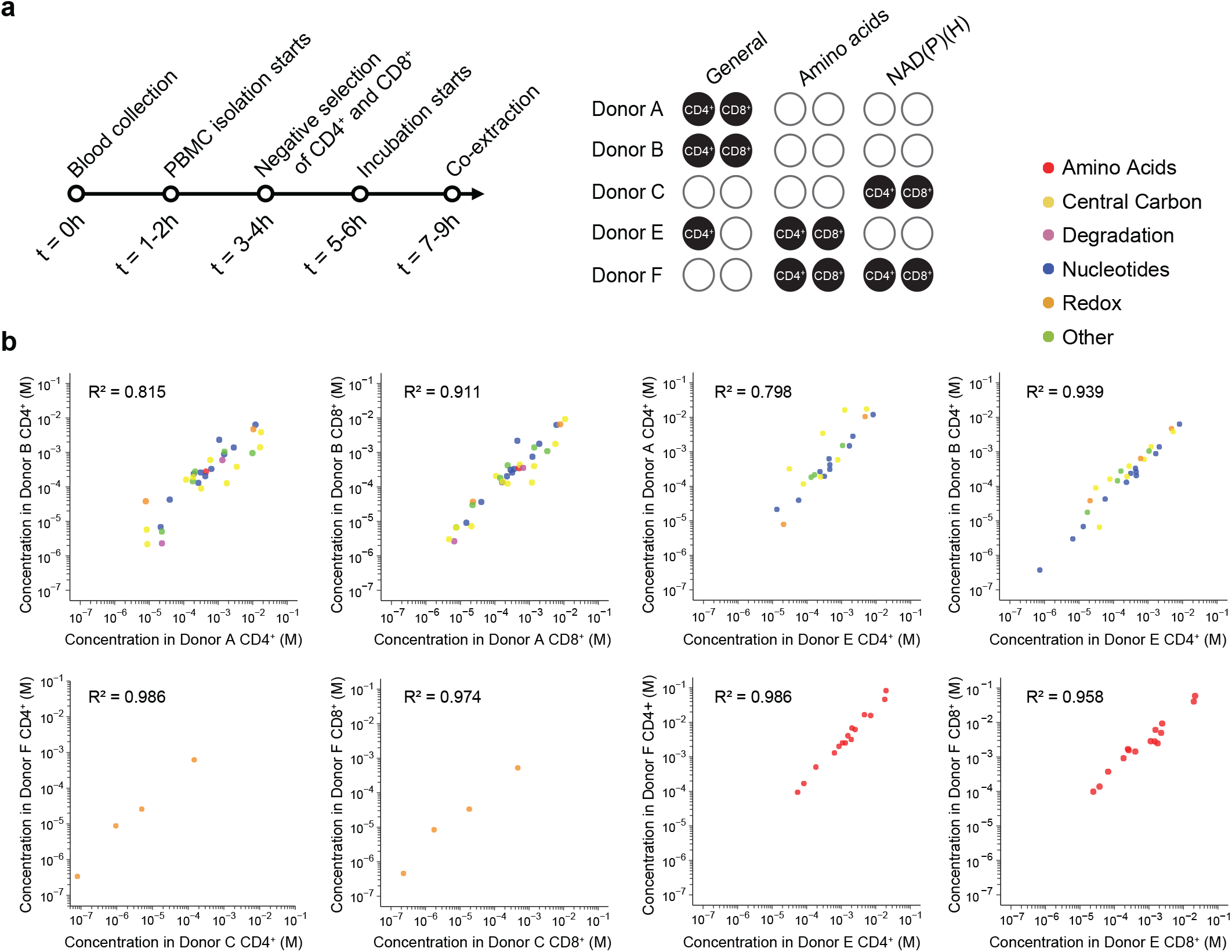
T cell metabolite concentrations are conserved across individuals. (**a**) Primary T cell metabolites were quenched and extracted within 7-9 hours of blood collection from five healthy donors. Different metabolite classes, which required different extraction methods, were measured for each donor because there were not enough T cells to measure all metabolite classes from any one individual. General metabolites included those in central carbon metabolism, degradation pathways, nucleotides and their biosynthesis pathways as well as some redox metabolites (e.g., glutathione) and others. (**b**) Absolute metabolite concentrations were quantified in CD4^+^ helper T cells and CD8^+^ killer T cells and were compared across donors.

We quantified 70 metabolites in CD4^+^ and CD8^+^ T cells from five donors (**Fig. 2b and Supplementary Tables 1 and 2**). T cell metabolomes were highly correlated across the five donors tested (R^2^>0.79) (**Fig. 2b**). While we did not obtain enough cells to quantify all 70 metabolites from all individual donors, the resemblance across individuals revealed their integral parts in a consolidated picture of human T cell metabolome. To account for individual variability and measurement uncertainty, we used weighted bootstrapping (see **Methods**) for unifying metabolite measurements from five donors (**Fig. 3a and Extended Data Fig. 2**). The total concentrations of measured metabolites spanning central carbon metabolites, amino acids, nucleotides, redox metabolites, and others in CD4^+^ and CD8^+^ T cells were 154 mM and 131 mM.

**Figure 3.**
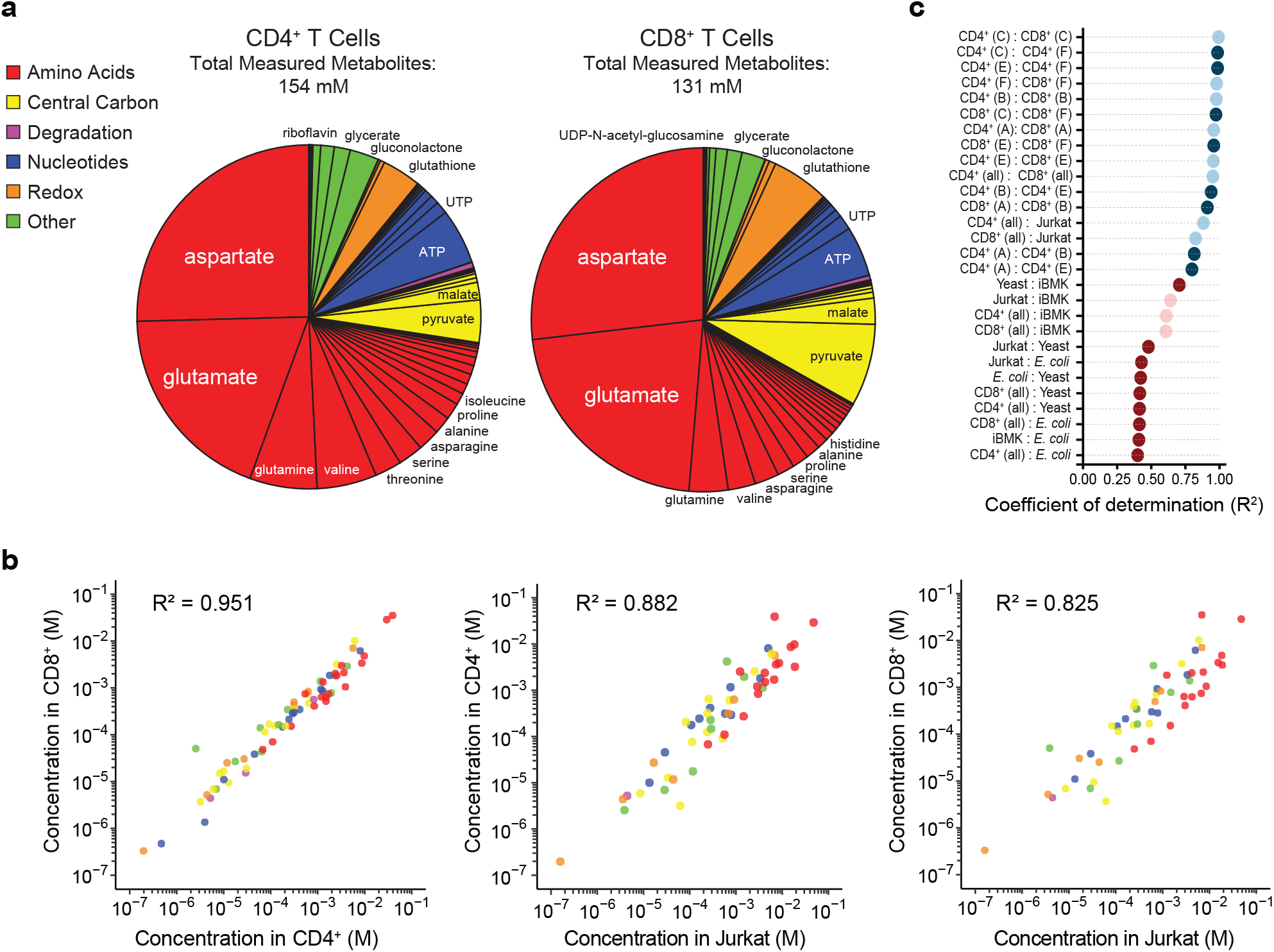
T cell metabolite concentrations are conserved across subtypes. (**a**) Absolute metabolite concentrations in primary T cells were unified in accordance with measurements from all experiments and individuals by bootstrapping (see **Methods**). (**b**) Metabolite concentrations were compared between primary CD4^+^ and CD8^+^ T cells and between primary and Jurkat T cells. (**c**) The metabolomes of T cell subtypes from different individuals, *E. coli*, yeast, iBMK, and Jurkat cells were compared. Light blue dots represent comparisons between T cell subtypes; dark blue dots between individuals; light red dots, between T cells and other mammalian cells; and dark red dots, between T cells and microorganisms.

Amino acids represented the most abundant metabolite group comprising 73%, 67%, and 67% of the measured metabolome in CD4^+^, CD8^+^, and Jurkat cells, respectively. The most abundant metabolite measured in primary T cells was aspartate whereas in Jurkat cells glutamate was the most abundant. Central carbon metabolites and nucleotides were the next most abundant groups, each comprising as much as ∼10% of the metabolome in primary T cells. Pyruvate and malate were consistently the most abundant central carbon metabolites, and ATP and UTP were the most abundant nucleotides. Interestingly, 3-phosphoserine and uridine were two of the most abundant metabolites measured in Jurkat cells, pointing to the role of *de novo* serine^28^ and pyrimidine^29^ biosynthesis in T cell proliferation and development.

### Conservation of T cell metabolome across subtypes

In addition to the broad resemblance between the metabolomes of T cell subtypes, we observed a near-perfect correlation (R^2^>0.95) between individual metabolite concentrations in CD4^+^ and CD8^+^ T cells (**Fig. 3b** and **Extended Data Fig. 3**). Despite some differences in some of the most abundant metabolites, as a whole, metabolite concentrations in Jurkat cells were highly correlated with those of CD4^+^ and CD8^+^ primary T cells (R^2^=0.88 and R^2^=0.83). To assess if these correlations were noteworthy, we compared the metabolomes of T cells to those of other systems (previously measured using the traditional isotope ratio-based approach)^23,24^. Across subtypes and individuals, all correlations of T cell metabolite concentrations had R^2^ values of 0.8 or greater, exceeding those of other comparisons whose R^2^ values ranged from 0.4 to 0.7 (**Fig. 3c**).

The lowest observed correlation was between T cells and *E. coli* with R^2^ around 0.4 (**Extended Data Fig. 4a**). Most amino acid concentrations were higher in T cells, whereas the concentrations of select central carbon metabolites, nucleotides, and redox cofactors were higher in *E. coli*. The T cell metabolome showed a higher correlation (R^2^>0.4) with yeast (*S. cerevisiae*) cells than with *E. coli* (**Extended Data Fig. 4b**). Both microbes possessed greater concentrations of glycolytic intermediates such as hexose phosphates, fructose-1,6-bisphosphate, and 3-phosphoglycerate. Metabolite concentrations in T cells were moderately correlated (R^2^>0.6) with those of iBMK cells (**Extended Data Fig. 4c**). On one hand, the observed R^2^ values across the kingdoms of life indicated a baseline conservation of metabolism beyond just its network structure. Between 40% and 65% of the variance in metabolite concentrations in human T cells could be accounted for by metabolite concentrations in disparate systems. On the other hand, the extraordinary conservation of the T cell metabolome across subtypes and individuals implied the existence of underlying design principles unique to T cell metabolism.

### T cell metabolome conforms to Zipf’s law

A recurring observation in metabolite profiles was the high abundance of select metabolites such as aspartate, glutamate, pyruvate, and ATP. Why do a few metabolites dominate the metabolome? To answer this question, we investigated whether metabolite concentrations followed Zipf’s law (**Eqn. 3**), which states that, when measured values are sorted in decreasing order, the n^th^-ranked measurement f(n) is approximately inversely proportional to n.

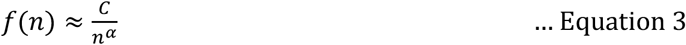

where α and C represent fitted parameters that are approximately 1 and the highest (n=1) measured value, respectively.

We overlaid Zipf’s law to ranked absolute metabolite concentrations such the fitted line goes through as many of the 95% confidence intervals of metabolite measurements as possible (see **Methods**). The top 35, 32, and 26 metabolites in CD4^+^, CD8^+^, and Jurkat cells followed Zipf’s law (**Fig. 4a**). Zipf’s law appears in many natural and artificial constructs (e.g., languages, city populations, and antibody nongenomic diversity^30^); however, the partial conformity of T cell metabolome to Zipf’s law was news and suggestive of the completeness of our measurement of high abundance metabolites and the efficiency of the uneven distribution of the metabolite pool.

**Figure 4.**
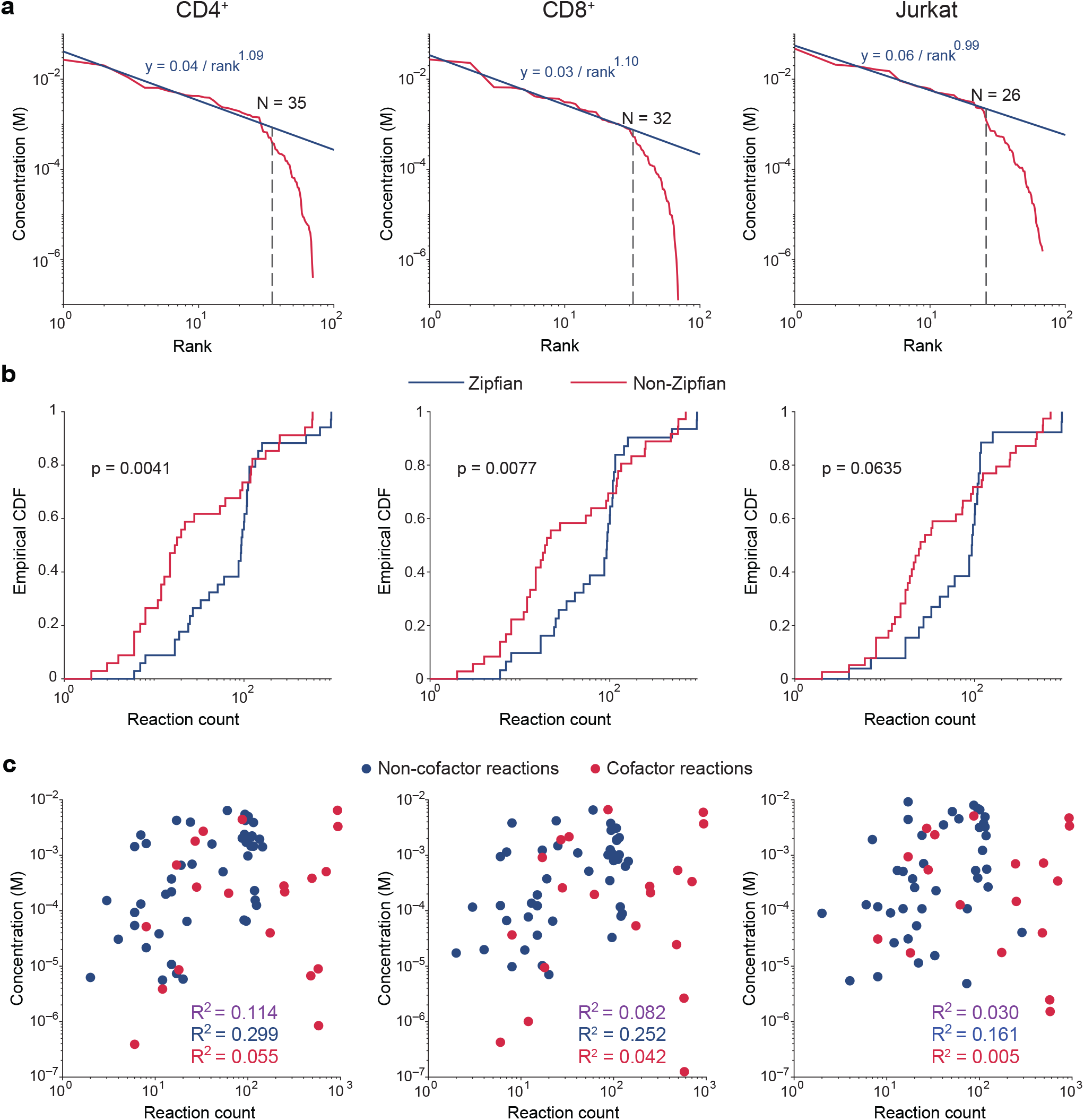
Top metabolites follow Zipf’s law and show high connectivity in metabolic networks. (**a**) Metabolite concentrations were rank-ordered from the highest to the lowest, and a line in accordance with Zipf’s law was fitted so that it goes through the maximum numbers (N) of metabolites’ 95% confidence intervals as possible. (**b**) The cumulative distribution function of total reaction counts associated with individual Zipfian (blue, rank≤N) and non-Zipfian (red, rank>N) metabolites. P values were computed from the Kolmogorov-Smirnov test comparing the two distributions within each T cell type. (**c**) The relationship between metabolite concentrations and connectivities (i.e., the numbers of reactions that individual metabolites participate in) was examined for non-cofactor metabolites (blue), cofactors (red), and all measured metabolites (blue+red=purple).

While the total metabolome pool size is bound by osmotics, the factors that affect the individual metabolite concentrations include metabolic network topology, reaction kinetics, and thermodynamics. We surmised that a metabolic network may operate more efficiently by concentrating metabolites positioned at highly interconnected nodes. Using a genome-scale reconstruction of human metabolism^31^, we counted the number of metabolic reactions each metabolite is acting as a substrate or a product. Zipfian metabolites whose concentrations were high participated in more reactions than non-Zipfian metabolites with low concentrations. (**Fig. 4b**). While global cofactors such as ATP, NADH, and NADPH participated in numerous reactions regardless of their concentrations being high or low, we observed a modest correlation (R^2^ as high as 0.3) between the connectivities and concentrations of non-cofactors (**Fig. 4c**). Aside from a few cofactors (e.g., ATP and CoA), T cells did not keep large pools of cofactors, which require onerous biosynthetic pathways, but instead recycled them since cofactors are not destroyed in the reactions that they facilitate. Thus, metabolic network topology corroborated metabolite concentrations supporting efficiency. While reaction kinetics and thermodynamics also shape the metabolome, a critical determinant of a metabolite’s concentration is its relevance to regulating biological functions beyond metabolism.

Aspartate, the most abundant metabolite in human primary T cells, is a central regulator of immune function^32-37^.

### T cells efficiently use and regulate enzymes

Enzyme-catalyzed reactions display saturable kinetics with respect to metabolite concentrations (i.e., Michaelis-Menten kinetics). Non-dimensionalized concentration [met]/K_m_, where [met] is the concentration of a metabolite and K_m_ is the Michaelis constant for an enzyme active site with respect to the metabolite, represents the occupancy of an enzyme active site: [met]/K_m_ ≥ 1 represents high occupancy and efficient enzyme use whereas [met]/K_m_ < 1 represents low occupancy and inefficient enzyme use. We observed that the distribution of non-dimensionalized concentrations (with true K_m_) was significantly narrower than the that with permutated K_m_ values as quantified by their interquartile ranges (IQRs) (**Fig. 5a**). The narrow distribution of non-dimensionalized concentrations suggested coevolution of enzymes and metabolite concentrations toward efficient enzyme use.

**Figure 5.**
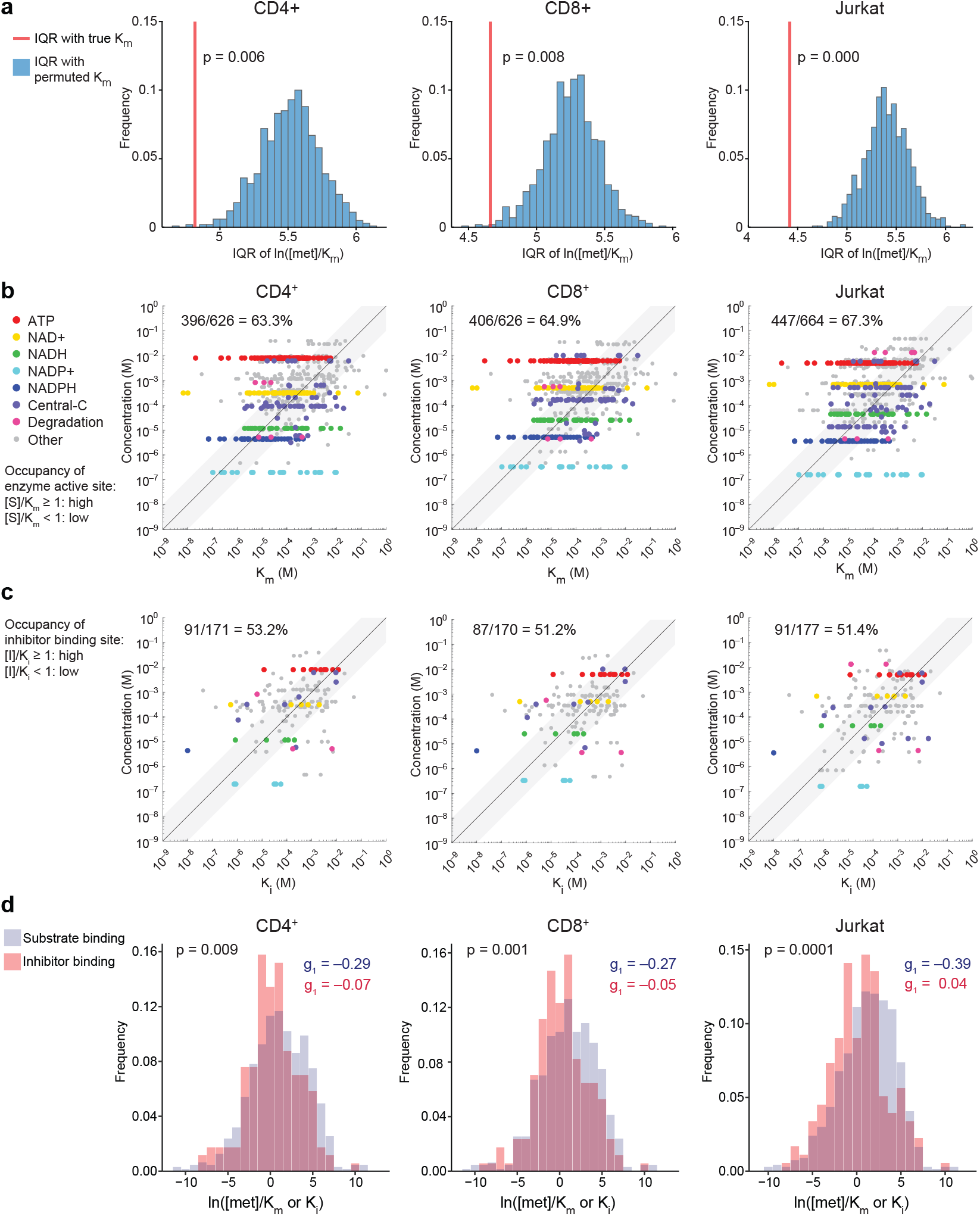
Absolute metabolite concentrations reveal efficient enzyme use. (**a**) The interquartile range (IQR) of ln([met]/K_m_) (red line) is significantly lower than that of ln([met]/K_m,permuted_), in which K_m_ values for enzyme active sites were randomly permuted. P value was twice the fraction of the IQRs with permuted K_m_ that were less than or equal to the IQR with true K_m_. (**b**) Absolute metabolite concentrations were compared with K_m_ for the associated enzyme active sites. The fraction of concentrations exceeding K_m_ is shown. (**c**) Absolute metabolite concentrations were compared with inhibitor dissociation constants (K_i_) for the associated inhibitor binding sites. The fraction of concentrations exceeding K_i_ is shown. (**d**) Tendencies for substrate-enzyme binding and inhibitor-enzyme binding were compared across T cell types. P values were from the Kolmogorov-Smirnov test comparing the distributions of ln([met]/K_m_) and ln([met]/K_i_). The g_1_ values represent the Fisher-Pearson moment coefficient of skewness.

Metabolites serve as both substrates and effectors for enzymes. As the substrate and the effector increasingly occupy the active site and the effector binding site of an enzyme, their influences on the reaction rate contract. Across all three T cell types, two thirds of substrate-enzyme pairs displayed metabolite concentrations exceeding their corresponding K_m_ (**Fig. 5b and Supplementary Tables 3, 4, and 5**). On the other hand, metabolites as inhibitors had their concentrations exceed the corresponding dissociation constants of the enzyme-inhibitor complex (K_i_) for half of inhibitor binding interactions (**Fig. 5c and Supplementary Tables 6, 7, and 8**). Accordingly, the distributions of log ratios of metabolite concentrations to corresponding K_m_ values were left-skewed, whereas log ratios of metabolite concentrations to corresponding K_i_ values appeared more symmetrical (**Extended Data Fig. 5**). In each of the three T cell types, the former distribution of ln([met]/K_m_) leaned to the right compared to the latter distribution of ln([met]/K_i_) (**Fig. 5d and Supplementary Table 9**). On the individual metabolite level, ATP and glutathione saturated the majority of enzyme binding sites as either substrates or inhibitors, whereas cyclic AMP and NADP^+^ hardly occupied enzyme binding sites (**Extended Data Fig. 6**). Non-saturating metabolites like cyclic AMP and NADP^+^ influence reaction rates through mass action. On the other hand, saturating metabolites like ATP and glutathione would have to decrease their concentrations drastically to meaningfully influence reaction rates. Nonetheless, they may still influence reaction rates by modulating ATP/ADP and GSH/GSSG ratios and shifting ΔG of near-equilibrium reactions^21,38^. The concentrations of central carbon metabolites and purine nucleotides (excluding ATP) were distributed more evenly around the respective K_m_ and K_i_ values, sensitizing reaction rates to changes in metabolite levels (presumably to promote adaptation). Collectively, absolute metabolite concentrations manifested efficient utilization of enzymes and modulation of enzyme activity.

### T cells maintain a high energy charge

Absolute metabolite concentrations conferred energetic and thermodynamic insights into T cell metabolism. ATP is the cellular energy currency. Exergonic ATP hydrolysis drives reactions, which are otherwise thermodynamically unfavorable, and produces ADP in the process. Thus, maintaining a high ATP-to-ADP ratio favors proper cellular function. In both primary and immortalized T cells, ATP/ADP ratios were consistently high with an IQR of 16.8–20.3 (**Fig. 6a**), which is higher than those of other eukaryotes yeast and iBMK cells (∼3.95 and ∼8.21, respectively)^24^. The high ATP/ADP ratio in resting T cells may prime them for a rapid switch to proliferation and cytokine synthesis upon activation. To fully account for cellular energetics, we quantified the adenylate energy charge (**Eqn. 4**), which reflects the total energy stored in the adenylate pool.

**Figure 6.**
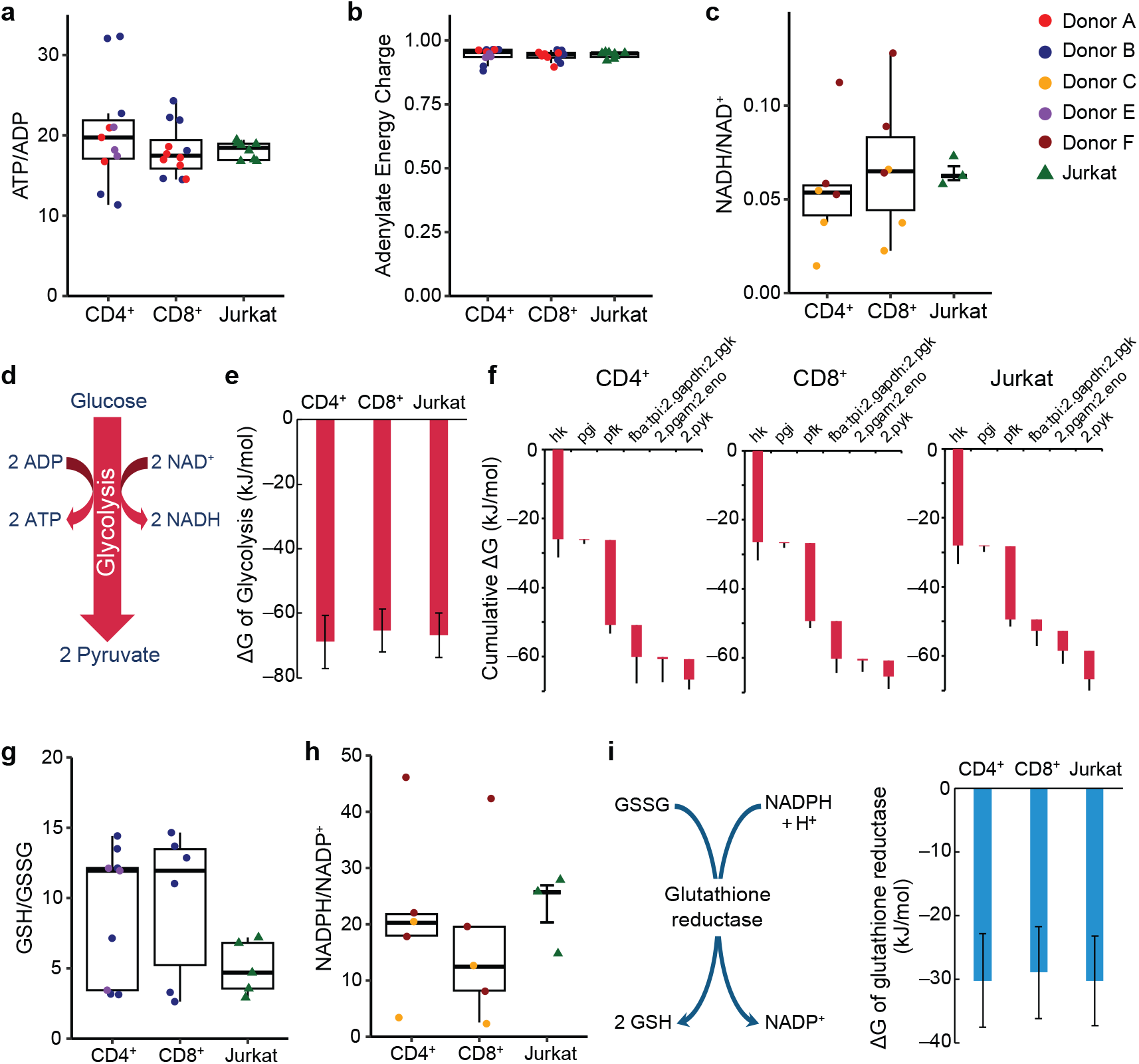
Energetics of T cell metabolism. (**a**) The ratio of ATP to ADP concentrations was calculated for primary CD4^+^ and CD8^+^ T cells and Jurkat cells. (**b**) The adenylate energy charge was calculated using the concentrations of ATP, ADP, and AMP. n=11 replicates derived from three donors for CD4^+^; n=12 replicates derived from two donors for CD8^+^ T cells; and n=8 replicates for Jurkat cells. (**c**) The ratio of NADH to NAD^+^ was calculated. n=6 replicates derived from two donors for CD4^+^ and CD8^+^ T cells; and n=3 replicates for Jurkat cells. (**d**) Glycolysis converts each molecule of glucose into two molecules of pyruvate while generating two ATP and two NADH. (**e**) Overall Gibbs free energy change across glycolysis in T cells was computed using the net reaction of glycolysis. (**f**) Using the absolute concentrations of glycolytic intermediates, stepwise and cumulative Gibbs free energy changes across glycolysis were computed. (**g**) The ratio of glutathione (GSH) to glutathione disulfide (GSSG) was calculated. n=9 replicates derived from two donors for CD4^+^ T cells; n=6 replicates derived from a single donor for CD8^+^ T cells; and n=5 replicates for Jurkat cells. (**h**) The ratio of NADPH to NADP^+^ was calculated. n=5 replicates derived from two donors for CD4^+^ and CD8^+^ T cells; and n=3 replicates for Jurkat cells. (**i**) Glutathione reductase generates reduced glutathione using the reducing power of NADPH. Gibbs free energy of glutathione reductase reaction was computed using substrate and product concentrations. Error bars represent the 95% confidence intervals. Boxplots show the quartiles with thick lines representing the median. The whiskers extend to the farthest measurement values within 1.5 times the IQR from the edges of the box.

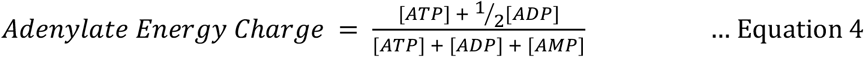

The adenylate energy charge was 0.93-0.96, tightly conserved across all T cell types (**Fig. 6b**). The observed adenylate energy charge in T cells was at the very high end of the physiological range from 0.7 to 0.96 in mammalian cells^24,39^.

### T cells maintain favorable redox ratios

Oxidative phosphorylation is the main source of ATP for aerobic organisms including humans. Using the proton motive force generated by electron transport chain from electron donors such as NADH and FADH_2_ to an electron acceptor O_2_, oxidative phosphorylation drives ATP synthesis. The NADH/NAD^+^ ratio in T cells ranged from 0.04 to 0.08 (IQR) (**Fig. 6c**). The ATP/ADP and NADH/NAD^+^ ratios affect the thermodynamic driving force of glycolysis, which generates two ATP and two NADH molecules for each glucose molecule (**Fig. 6d**). We determined the Gibbs free energy change (ΔG) across glycolysis to range from –60 kJ/mol to –77 kJ/mol (**Fig. 6e and Supplementary Table 10**). Three quarters of Gibbs free energy drop occurred in upper glycolysis (**Fig. 6f**). Thus, the low NADH/NAD^+^ ratio played dual roles of driving glycolysis forward (despite the high ATP/ADP) and preventing reactive oxygen species (ROS) production.

Redox pairs also play a vital role of providing reducing power for scavenging ROS and anabolism. Glutathione and NADPH are primary reducing agents in the cell. In T cells, glutathione plays a crucial role in effector function^40^. T cells maintained a high ratio of antioxidant glutathione (GSH) to its oxidized counterpart glutathione disulfide (GSSG) with the GSH/GSSG IQR ranging from 4 to 14 (**Fig. 6g**), which was in line with healthy erythrocytes whose GSH/GSSG=5.88 (ref. ^41^) but not as high as those observed in human tissues^24,42^ and whole blood^43^. For NADPH/NADP^+^, we observed large variations, but they were mostly high between 9 and 27 in T cells (**Fig. 6h**). Such variability was consistent with prior observations of NADPH/NADP^+^ of 2.30 in iBMK cells^24^ and ∼30 in immortalized mouse embryonic fibroblasts^44^. In addition to driving anabolism, the high NADPH/NADP^+^ ratio provided a driving force for the glutathione reductase reaction with ΔG= –30±7 kJ/mol (**Fig. 6i**). The conserved cellular energetics and thermodynamics across the three T cell types revealed T cells’ readiness for increased energy demand and oxidative stress.

## Discussion

Our knowledge of adaptive immunity builds on a strong foundation of cellular and molecular interactions between immune cells, receptors, cytokines, antigens, and antibodies as well as the ensuing signaling cascades. What powers these biochemical processes? How can immune cells invariably perform such critical tasks in varying environments? Does human immune diversity manifest in metabolism? The present work complements the growing body of knowledge in immunology by providing the first ever quantitative profiling of the human T cell metabolome in absolute terms.

The governing equations of life mathematically describe, at the local level, the kinetics and thermodynamics of biochemical reactions, and at the systems level, the laws of conservation of mass and energy. Absolute metabolite concentrations are integral elements of the governing equations. Since analytical instruments have varying sensitivities and responses to disparate analytes, however, absolute metabolome quantitation has remained an onerous task that requires large sample sizes. In particular, absolute metabolite concentrations in primary cells and tissues have been inaccessible due to low availability of biological materials. To solve this problem, we developed the ensemble method for quantifying absolute metabolite concentrations *en masse*.

With our ensemble quantitation method in hand, we quantified absolute metabolite concentrations in human T cells for the first time. T cell metabolism underlies adaptive immunity by supplying cellular energy and biochemical building blocks for activation, differentiation, and proliferation^7-10^ as well as production of cytokines and cytotoxic granules^16,45-48^. Thus, knowledge of absolute concentrations provides a quantitative and mechanistic understanding of T cell metabolism and establishes a solid foundation for promoting immunity^49,50^ and developing immunotherapy for cancer^14,51-53^ and autoimmune diseases^32,54-56^.

A key finding from our ensemble quantitation in human primary T cells is the conservation of metabolome across subtypes and individuals. Between five individuals and their helper (CD4^+^) and killer (CD8^+^) T cells, the lowest R^2^ observed was 0.8. Despite helper and killer T cells carrying out distinct functions in the adaptive immune system, their basal metabolic profiles resemble each other and indicate energy and redox homeostasis. On the one hand, the high metabolic similarity across individuals implies that metabolic variability may not stand in the way of allogeneic immunotherapy. On the other hand, ensemble absolute metabolome quantitation across a larger number of individuals may reveal the unique metabolic features of a “super donor,” whose cells would lead to successful treatment of many patients. Our data provides a baseline for healthy T cell metabolism, opening the door to biomarker discovery.

The observed conservation of metabolite concentrations manifests a metabolic design principle: cells prioritize concentration homeostasis despite a high turnover of metabolites. How can metabolites exert control of metabolic activity without a substantial change in concentrations? In reactions that are reversible or moderately forward driven, small changes in metabolite concentrations lead to large changes in reaction rates^21,57^. Furthermore, since each metabolite is involved in multiple reactions and pathways as substrates, products and effectors, a substantial change in a single metabolite’s concentration may have multiple unintended consequences throughout metabolism^58^. Therefore, small variations across multiple metabolites, rather than large variations of few metabolites, may better exert precise pathway-level modulation of metabolic activity. Caution should be exercised, however, not to misconstrue the overall similarity as lack of unique metabolic features.

The most notable difference between primary and immortalized T cells was the abundance of aspartate. In primary T cells, aspartate is the most abundant metabolite at 35-39 mM, whereas in Jurkat cells it just makes the top ten with 6.9 mM. Besides protein synthesis, aspartate serves three important functions: i) electron translocation from the cytosol to the mitochondrial matrix as part of the malate-aspartate shuttle; ii) biosynthesis of asparagine and nucleotides; and iii) anaplerosis. In T cells, both aspartate and asparagine have been linked to stress response pathways and effector function. Aspartate deficiency is a biomarker of autoimmune T cells from rheumatoid arthritis patients^32^. Aspartate deficiency lowers NAD^+^/NADH and ADP-ribosylation of GRP78/BiP, a regulator of endoplasmic reticulum (ER) stress, causing the expansion of ER membrane and the biogenesis of a proinflammatory cytokine tumor necrosis factor (TNF). Aspartate supplementation was sufficient to recover healthy T cell function. Asparagine availability and asparagine synthetase (ASNS) activity modulate T cell metabolism, effector function, and antitumor response^59-61^. The immunometabolite itaconate decreases CD8^+^ T cell proliferation in part by reducing aspartate, glycine and serine biosynthesis flux in CD8^+^ T cells^33^. Supplementation with aspartate, glycine and serine or nucleosides restores T cell proliferation, with aspartate supplementation being essential for the anti-tumor effector function of CD8^+^ T cells. Thus, aspartate is a key regulator of T cell inflammatory response (*cf*. succinate and itaconate in macrophages^62-64^) and we surmise that a high aspartate concentration is a hallmark of healthy T cells.

In addition to uncovering the conserved features of T cell metabolism, a key contribution of this work is the ensemble method for absolute metabolome quantitation, which standardizes metabolomics, augments its utility, and engenders kinetic and thermodynamic insights. The ensemble method can be applied to not only suspension cultures but also adherent cell cultures and tissues (**Extended Data Fig. 7**). Knowledge of absolute metabolite concentrations transcends the boundaries of individual metabolomics studies, which cannot be unified due to batch effects, and lays the foundation for cumulative knowledgebase of metabolism. It facilitates the determination of fluxes in non-steady-state conditions^65,66^, quantification of organelle-specific metabolites^67^, identification of metabolite-protein interactions^68,69^, and integration of multi-omic datasets^70^. We envision our ensemble quantitation method being applied to the >400 major cell types in the human body^71^ to create an atlas of human metabolome. The resulting knowledge will serve as a conduit for connecting biochemistry to mathematical modeling, advance predictive biology, and reveal master regulators of metabolism.

## Methods

### Negative selection of CD4^+^ and CD8^+^ T cells

CD4^+^ and CD8^+^ primary T cells were isolated from peripheral blood mononuclear cells (PBMCs) of healthy donors by magnetic-activated cell sorting (MACS). PBMCs were isolated from healthy donor blood (courtesy of the UCLA Blood and Platelet Center) by density gradient centrifugation using Ficoll-Paque PLUS (Cytiva) (courtesy of the UCLA Virology Core Laboratory). To prevent unintended metabolic reprogramming due to the binding of antibodies to the surface antigens of T cells^14,72^, negative selection strategies for CD4^+^ and CD8^+^ were employed. Manufacturer protocols for CD4^+^ and CD8^+^ negative selection kits (Miltenyi Biotec) were followed. PBMCs were spun down, rinsed with PBS, and resuspended in PBS containing 0.5% BSA and 2 mM EDTA (Miltenyi Biotec) at a concentration of 40 μL buffer per 10^7^ cells. Biotin-conjugated antibody cocktail (Miltenyi Biotec) was added at 10 μL per 10^7^ cells, and the cells were incubated for 5 minutes in the refrigerator (2-8°C). After allowing antibodies to bind to cognate antigens, 30 μL of buffer per 10^7^ cells was added along with 20 μL of MicroBead Cocktail (Miltenyi Biotec) per 10^7^, and the cells were incubated for 10 minutes in the refrigerator (2-8°C). For negative selection of CD4^+^ T cells, biotin-conjugated antibody cocktail included antibodies against CD8, CD14, CD15, CD16, CD19, CD36, CD56, CD123, TCR γ/δ, and CD235a (Glycophorin A), and MicroBead Cocktail included MicroBeads conjugated to anti-biotin antibody and anti-CD61 antibody to remove CD8^+^ T cells, monocytes, neutrophils, eosinophils, B cells, dendritic cells, NK cells, granulocytes, γ/δ T cells, erythroid cells, megakaryocytes, and platelets. For negative selection of CD8^+^ T cells, biotin-conjugated antibody cocktail included antibodies against CD4, CD15, CD16, CD19, CD34, CD36, CD56, CD123, TCRγ/δ, and CD235a (Glycophorin A), and MicroBead Cocktail included MicroBeads conjugated to anti-biotin antibody, anti-CD14 antibody, and anti-CD61 antibody to remove CD4^+^ T cells, monocytes, neutrophils, eosinophils, B cells, stem cells, dendritic cells, NK cells, granulocytes, γ/δ T cells, erythroid cells, megakaryocytes, and platelets.

LS columns (Miltenyi Biotec) were rinsed with buffer, and then the cells were added to the columns. The initial flow through represented the enriched CD4^+^ or CD8^+^ population. Once isolated, the cells were spun down at 200g for 10 minutes, the supernatant was decanted, and the cells were resuspended in RPMI supplemented with 10% dialyzed fetal bovine serum (dFBS). CD4^+^ or CD8^+^ T cells were incubated at 37°C and 5% CO_2_ for 1-2 hours before extraction. For amino acid measurement, cells from Donor E and Donor F were cultured for two hours in standard RPMI 1640 media diluted threefold with amino acid deprived RPMI media supplemented with 10% dFBS.

Population purity was verified with flow cytometry (**Extended Data Fig. 8**). To confirm the purity of isolated CD4^+^ and CD8^+^ T cells from PBMCs, cells were stained with the following antibodies: CD3 (APC, clone REA613, Miltenyi Biotec), CD4 (VioBlue, clone REA623, Miltenyi Biotec), and CD8 (FITC, clone REA715, Miltenyi Biotec). A suspension of 5×10^5^ cells in 100 μL of PBS containing 0.5% BSA and 2 mM EDTA was prepared. After 2 μL of each antibody solution was added, the cells were incubated on a shaker at room temperature in the dark for 10 minutes, washed with 1 mL of buffer, centrifuged at 300g for 5 minutes, and resuspended to a volume of 0.5 mL of staining buffer. Samples were moved to a flow cytometry tube and stored cold (2-8°C) in the dark until transport. Unstained and fluorescence minus one (FMO) control samples were prepared similarly. Single color controls were prepared following the same protocol with positive and negative compensation beads (Miltenyi Biotec) with the same concentration of antibodies added (2 μL antibody solution per 100 μL PBS containing 0.5% BSA and 2 mM EDTA. Cells were analyzed using an LSRFortessa flow cytometer (BD Biosciences), and data were gated and analyzed with Floreada.io.

### Cell lines and culture conditions

All cells were cultured in nutrient replete conditions. Jurkat cells (Jurkat clone E6-1, ATCC) were cultured in RPMI 1640 (Gibco) supplemented with 10% dFBS at 37°C and 5% CO_2_. For Jurkat cells, partial media change, replacing one third of the cell-free culture volume, was performed every 24 hours and three hours prior to metabolite extraction. Primary T cells were seeded into RPMI 1640 (Gibco) supplemented with 10% dFBS at 37°C after negative selection, and their intracellular metabolites were extracted within one to two hours. The cells designated for intracellular amino acid quantification and from Donor E were seeded into a medium with one-third diluted amino acids but otherwise identical to RPMI 1640 supplemented with 10% dFBS. *E. coli* strain NCM3722 was grown in Gutnick minimal medium^73^ containing 2 g/L [U-^13^C_6_] glucose and 10 mM NH_4_Cl at 37°C. *E. coli* cells were cultured in the ^13^C-labeled medium starting from the inoculation of overnight cultures. Cells were reinoculated from the overnight culture into beveled flasks and grown for four to five hours to the mid-exponential phase before metabolite extraction. *BAX*^−/−^/*BAK*^−/−^ immortalized baby mouse kidney (iBMK) cells^74^ cells were cultured in DMEM without glucose, pyruvate, and glutamine (Gibco) supplemented with 4.5 g/L [U-^13^C_6_] glucose, 584 mg/L [U-^13^C_5_] glutamine (99%, Cambridge Isotope Laboratories) and 10% dFBS in six-centimeter plates at 37°C and 5% CO_2_ for 72 hours. Fresh medium was replenished every 24 hours and three hours (2 g/L [U-^13^C_6_] glucose but otherwise the same medium) prior to metabolite extraction.

### Co-extraction of metabolites from two cell types

Jurkat, *E. coli*, and iBMK cells were all extracted when they were exponentially growing. Jurkat cells were extracted when culture density was between 1×10^6^ cells/mL to 3×10^6^ cells/mL. *E. coli* cells were extracted in the mid-exponential phase at OD_600_ between 0.3 and 0.6. iBMK cells were extracted when cultures reached 60-80% confluence. Primary T cells were extracted one to two hours after isolation from PBMCs, after incubating in RPMI 1640 + 10% dFBS. Viability and growth of mammalian cells were measured by mixing cell suspension with Trypan Blue at a 1:1 ratio and using the Countess II FL Cell Counter (Thermo).

For co-extraction of T cells and *E. coli* cells, the two cultures were simultaneously vacuum filtered by pipetting them onto distinct regions of a 47-mm nylon membrane (0.20-µm pore size). Care was taken to ensure that the two cultures did not come in contact. We employed three quenching-extraction methods tailored to redox cofactors, metabolites present in the extracellular space (e.g., amino acids and pyruvate), and all others including central carbon metabolites (**Extended Data Fig. 9a**). Once the cells were separated from the media, the filter was placed cell-side down onto a 6-cm plate containing 1 mL of pre-cooled (– 20°C) 40:40:20 LC-MS grade methanol/acetonitrile/water. Extraction continued in a –20°C freezer for 15-20 minutes. The filter was flipped so that the cell side was facing up. The extract solution was mixed well, transferred to an Eppendorf tube, and spun down at 17,000g and 4°C for 10 minutes. For measuring NAD(P)(H), an acidic extraction solvent containing 0.1 M formic acid was used^75^ to quench residual enzymatic activity and dissociates metabolites by fully denaturing enzymes, and prior to centrifugation, the extract was neutralized with 15% NH_4_HCO_3_ (**Extended Data Fig. 9b**). The supernatant was dried under nitrogen gas and reconstituted in LC-MS-grade water for LC-MS analysis. The formic acid extraction enabled the quantification of NADPH and NADH in a reproducible manner (**Extended Data Fig. 9c**).

Measurement of intracellular amino acids in T cells required rapidly washing cells to minimize the perturbation of cellular metabolism. Initially, two washing methods, in which 150 mM ammonium acetate at pH 7.4 or PBS (Corning) was pipetted onto cells on nylon filter immediately after vacuum filtration, were tested; however, both led to cell lysis and loss of intracellular metabolites (**Extended Data Fig. 9d**). Instead, a combination of centrifugation and vacuum filtration was used for rapid washing and extraction. T cells were first separated from media by centrifuging 1 mL of culture at 500g for 2 minutes. The supernatant was removed, and cells were resuspended in 1 mL of amino acid-free RPMI 1640 before rapid vacuum filtration and co-extraction as described above (**Extended Data Fig. 9e**). One wash step was sufficient to quickly remove extracellular amino acids (**Extended Data Fig. 9f**). For Jurkat cells, which secreted substantial amounts of pyruvate, pyruvate measurement also followed rapid washing and extraction steps as with amino acids.

For co-extraction of T cells and iBMK cells, both culture plates were removed from the incubator simultaneously. Media was aspirated from the iBMK culture plate, extraction solvent was placed on it, the plate was placed on a cold plate. Jurkat cell culture was vacuum filtered on a nylon filter membrane, and the filter was placed onto the iBMK extraction plate with the cell side facing down onto the extraction solvent (**Extended Data Fig. 7a**). The plate was placed in the –20°C freezer for 15-20 minutes. The adherent cells were detached using a cell lifter, the filter was flipped cell-side up, and the extract solution was mixed well. The metabolite extract was collected, spun down at 17,000g and 4°C for 10 minutes. The supernatant was dried under nitrogen gas and reconstituted in LC-MS-grade water for LC-MS analysis.

### Metabolite measurement

The metabolite extracts were analyzed by LC-MS, a Vanquish Duo UHPLC (Thermo) coupled to a Q Exactive Plus mass spectrometer (Thermo). The LC separation was achieved using a hydrophilic interaction chromatography column (XBridge BEH Amide Column, 130 Å, 2.5 µm, 2.1 mm × 150 mm, Waters). The A and B solvents were 95:5 water/acetonitrile with 20 mM ammonium acetate, 39 mM ammonium hydroxide (pH 9.4) and 99:1 acetonitrile/water, and the gradient was the same as previously reported^14,76^. MS was performed in positive and negative ion mode with a mass resolution of 140,000 at 200 *m/z* (mass-to-charge ratios). LC-MS data were processed using the Metabolomic Analysis and Visualization Engine (MAVEN)^77^. The m/z values from monoisotopic masses and the retention times from authenticated standards were used to identify peaks. For isomers glucose-6-phosphate and fructose-6-phosphate, which were not separated by the LC method, the ratio of isomers in the cells of interest was assumed to be the same as in the reference cells, and a single peak and U/L ratio were taken for both isomers. For arginine and glycine, whose concentrations could not be reliably obtained from co-extraction, known concentrations of arginine and glycine standards were spiked into cellular extracts, and cellular arginine and glycine concentrations were computed using the increase in the ion counts relative to the unspiked aliquots and the ion counts in the unspiked aliquots. A previously reported concentration value for dTTP concentration in *E. coli* was inaccurate. To measure the correct absolute concentration of dTTP in *E. coli, E. coli* strain NCM3722 was grown in Gutnick minimal medium^73^ containing 2 g/L [U-^13^C_6_] glucose and 10 mM NH_4_Cl at 37°C, and metabolites were extracted using 40:40:20 methanol/acetonitrile/water containing a known amount of unlabeled dTTP. The samples were prepared and analyzed as described above, and the new dTTP concentration was calculated from the ratio of unlabeled to labeled dTTP. The new dTTP concentration in *E. coli* was 1.81×10^-4^ M with the 95% confidence interval [1.54×10^-4^ M, 2.13×10^-4^ M]. Using this new reference concentration, dTTP concentrations in T cells were determined using the co-extraction method.

### Absolute metabolome quantitation and statistical analysis

Absolute metabolite concentrations were determined by the ratio of unlabeled to ^13^C-labeled metabolites, the ratio between aqueous cell volumes of cells of interest and reference cells, and the known absolute concentration in the reference cells. Measured U:L ratios depend on the intracellular aqueous volumes (V) and metabolite concentrations ([met]) in each cell type:

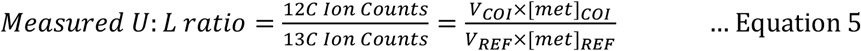

The concentration of each metabolite was calculated by:

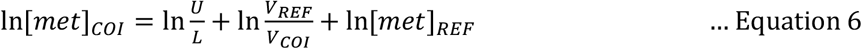

Where [*met*]_*COI*_ and [*met*]_*REF*_ is the metabolite concentration in the cells of interest (COI) and reference cells (REF), 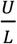 is the ratio of unlabeled ions to labeled ions, *V*_*COI*_ and *V*_*REF*_ are the aqueous cell volumes of COI and REF. For *E. coli*, culture density (OD_600_) was measured using a spectrophotometer and converted to aqueous cell volume using the conversion^24^: 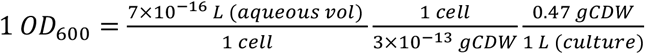. For primary T cells, Jurkat cells, and iBMK cells, packed cell volume was measured and converted to the aqueous cell volume using the conversion 1 *μL* (*PCV*) = 0.7 *μL* (*aq vol*).

Once metabolites concentrations were determined using **Eqn. 6** for individual samples, average levels and confidence intervals for metabolites were derived by bootstrapping. Since 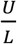 values influenced the accuracy of metabolite concentration measurement (**Fig. 1f**), bootstrapping was carried out by weighting metabolite concentrations based on 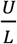. The weights were derived from the relationship between 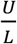 and the corresponding R^2^ values across the metabolome in the co-extraction validation study (**Fig. 1f**).

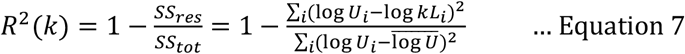

k represents expected 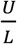 ratio (1, 4, and 20 in the validation study), i represents metabolite index, and U_i_ and L_i_ represent ion counts for unlabeled and ^13^C-labeled metabolite i.

The weights for individual metabolite concentrations in bootstrapping were determined by the likelihood of the measurements given their 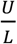 values accurately capturing the true concentrations, which were represented by R^2^. To determine weights for the individual samples whose metabolites were extracted from various 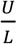 values, the known k and R^2^ values were fitted to a gaussian curve using nonlinear regression (**Extended Data Fig. 2**). The resulting weight function 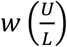 was used to determine resampling probabilities for weighted bootstrapping of T cells concentrations measurements.

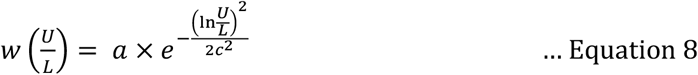

The values a = 0.831451 and c = 4.575862 were determined by non-linear least squares fitting. Each measured 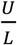 ratio was used to determine a metabolite concentration using **Eqn. 6** and a weight value *w*_*i*_ using **Eqn. 8**. For each metabolite, the probability of selecting a concentration measurement in weighted bootstrapping was computed by dividing its weight by the sum of the weights of all measurements:

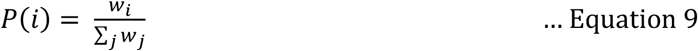

For each metabolite, the log-concentration (In[*met*]_*COI*_) distribution was bootstrapped by resampling 10,000 means, each of which was calculated from n (the number of samples) log-concentration measurements selected according to *P*(*i*) with replacement (**Extended Data Fig. 2**). The standard error *s*_*met*_ of log concentrations from the bootstrapped distribution was obtained by dividing the range of the 95% confidence interval by 4. However, this error represented a compounding effect of independent random errors from both COI (T cells) and REF cells. The errors for metabolites specifically in COI (T cells) were propagated using the compound standard error *s*_*met*_ and the standard error of 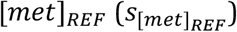:

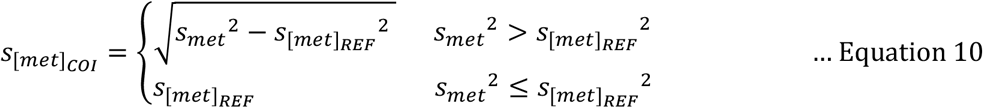

The 95% confidence interval of each metabolite was determined as the mean 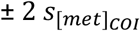 in logarithmic space:

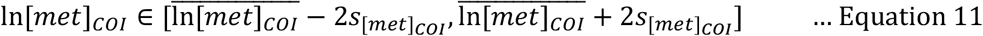

### Metabolome distribution analysis using Zipf’s law and genome-scale metabolic model

Metabolite concentrations were randomly sampled from the respective 95% confidence intervals and ranked from the largest to the smallest. Concentrations were log-transformed and used to fit log-transformed Zipf’s law:

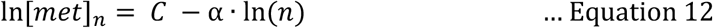

where n is the rank (with the highest concentration corresponding to n=1), [met]_n_ is the n^th^-ranked metabolite concentration, 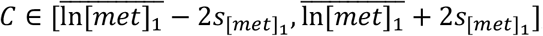, and *α* ∈ [0.90, 1.10]. Metabolite concentrations partially followed Zipf’s law. Thus, the number of Zipfian metabolites was determined as the largest rank N such that the best-fit line described by **Eqn. 12** for metabolites ranked 1 through N fell within the bounds set for C and *α*. The random sampling and fitting processes were repeated 10,000 times to find the maximum N. The instance shown in **Fig. 4a** was the first permutation encountered with the maximum N.

The numbers of reactions that individual metabolites are involved with were obtained from Recon3D, a genome-scale reconstruction of human metabolism^31^. For each measured metabolite, all rows of the S matrix of Recon3D corresponding to the metabolite were probed, and all non-zero columns therein corresponding to reactions (including transport reactions) were counted. This reaction count represented the total number of unique reactions across all compartments.

### Michaelis constants (K_m_) and inhibitor dissociation constants (K_i_)

Absolute metabolite concentrations were compared to known Michaelis constants (K_m_) and inhibitor dissociation constants (K_i_). K_m_ and K_i_ values in *Homo sapiens* were extracted from BRENDA using the Python 3 implementation of the Simple Object Access Protocol (SOAPpy)^78^. Entries corresponding to mutant enzymes were removed. When multiple entries for the same enzyme-metabolite pair were available, K_m_ or K_i_ was represented by the geometric mean. The true interquartile range of ln([met]/K_m_) was compared to a distribution of interquartile ranges determined by 1,000 random permutations of K_m_ values for enzyme active sites. The ln([met]/K_m_) and ln([met]/K_i_) distributions were compared using the Kolmogorov-Smirnov test. The skewness of each distribution was represented by the Fisher-Pearson coefficient of skewness (*g*_1_).

### Gibbs free energy of reaction

Gibbs free energies of lumped glycolytic steps and glutathione reductase were calculated using the eQuilibrator API^79^ by inputting the measured absolute concentrations of glycolytic intermediates, ATP, ADP, inorganic phosphate, NAD(H), NADP(H), glutathione, and glutathione disulfide. Inorganic phosphate concentration was set to the geometric mean of the range 2.5-5 mM based on the phosphate concentration in RPMI 1640 of 5.6 mM (Gibco) and previous measurements in mammalian cells^80,81^. Intracellular glucose concentration was set to the geometric mean of the range 0.1-6 mM based on previous measurements in mammalian cells^82-84^. For intracellular inorganic phosphate and glucose, the standard errors of their log concentrations were obtained by dividing the reported range in logarithmic space by 4.

Gibbs free energy changes for lumped reactions were calculated when intermediate metabolite concentrations were unknown. Gibbs free energy changes for the following reactions and lumped reactions were calculated.

Hexokinase (HK) reaction:

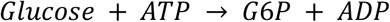

Phosphoglucoisomerase (PGI) reaction:

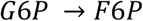

Phosphofructokinase (PFK) reaction:

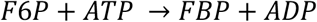

Lumped fructose-bisphosphate aldolase (FBA), triosephosphate isomerase (TPI), glyceraldehyde-3-phosphate dehydrogenase (GAPDH), phosphoglycerate kinase (PGK) reactions:

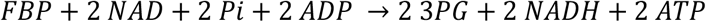

Lumped phosphoglycerate mutase (PGAM), enolase (ENO) reactions:

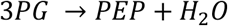

Pyruvate kinase (PYK) reaction:

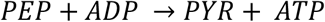

Overall glycolysis:

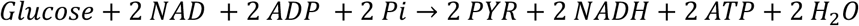

Glutathione reductase:

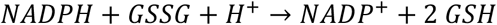

The error for each (lumped) reaction step was calculated by error propagation with the standard errors of the standard Gibbs free energy and the log concentrations of substrates and products (**Supplementary Table 10**). For instance, the error propagation formula for the Gibbs free energy of a reaction with the form *aA* + *bB* → *cC* + *dD*, where lowercase letters represent stoichiometric coefficients for metabolites in capital letters was:

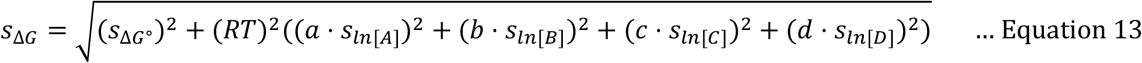

## Data availability

Source data for all Figures (absolute metabolite concentrations, kinetic constants, and thermodynamic constants) are provided in Supplementary Tables 1-10 and the GitHub public repository: https://github.com/samokeeffe/abs-met-quant

## Code availability

The code for bootstrapping and computational analyses is available on the GitHub public repository: https://github.com/samokeeffe/abs-met-quant

## Acknowledgements

The authors would like to thank the members of the Park lab, the UCLA Molecular Instrumentation Center, the UCLA Metabolomics Center, the UCLA-CDU Center for AIDS Research (CFAR), and the UCLA Flow Cytometry Core for helpful discussion. PBMCs were obtained from the UCLA-CDU CFAR Virology Core Lab that is supported by National Institutes of Health grant number 5P30 AI028697. Flow cytometry was performed in the UCLA Jonsson Comprehensive Cancer Center (JCCC) and Center for AIDS Research Flow Cytometry Core Facility that is supported by National Institutes of Health awards P30 CA016042 and 5P30 AI028697 and by the JCCC, the UCLA AIDS Institute, the David Geffen School of Medicine at UCLA, the UCLA Chancellor’s Office, and the UCLA Vice Chancellor’s Office of Research. This work was supported by the National Institutes of Health under instrumentation grant number 1S10OD016387-01, the National Center for Complementary and Integrative Health of the National Institutes of Health under award number R21AT012814 (J.O.P.), and the National Institute of General Medical Sciences of the National Institutes of Health under award number R35GM143127 (J.O.P.).

## Contributions

S.O.K. and J.O.P. designed the study and wrote the manuscript. S.O.K., H.S., E.M., and J.O.P. carried out the experiments. S.O.K. and H.S. isolated primary T cells and performed flowcytometry. S.O.K., H.S., E.M., and J.O.P. performed metabolomics analysis.

## Extended Data Figures

**Extended Data Figure 1.**
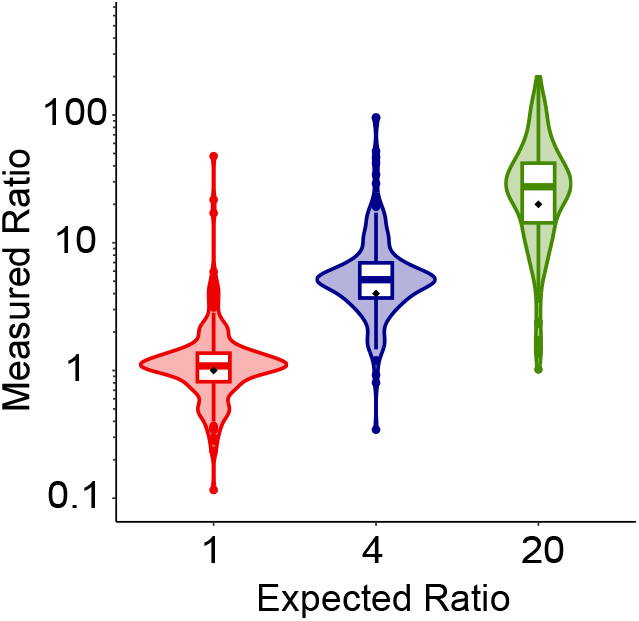
Co-extraction across ranges of metabolite ratios. The measured ratios of unlabeled (^12^C) to ^13^C-labeled metabolites from unlabeled and labeled *E. coli* cultures co-extracted at the ratios of 1:1, 4:1 and 20:1 were compared to the expected ratios (black diamonds).

**Extended Data Figure 2.**
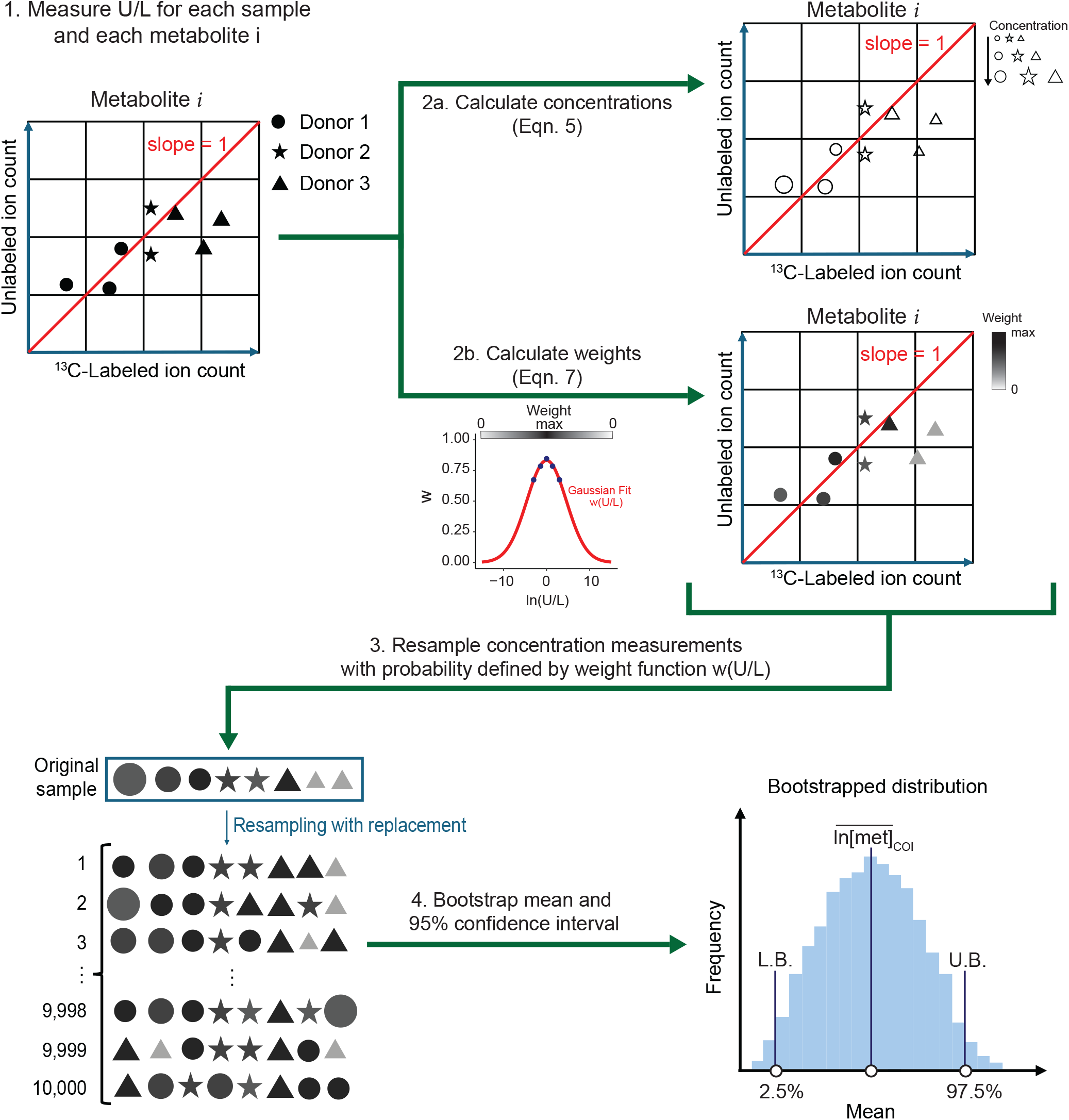
Determination of the distribution of log-concentrations of metabolites by bootstrapping. To unify metabolite measurements from multiple experiments and individuals and to determine the mean and 95% confidence interval of each metabolite concentration, a weighted bootstrapping approach was employed. The Gaussian fit w(U/L) assigned a weight for resampling each metabolite concentration measurement using its associated U/L value. Log-concentration distributions were bootstrapped by resampling 10,000 means, each of which was calculated from weighted sampling of n (the number of samples) log-concentration measurements with replacement. The upshot was a reference human T cell metabolome.

**Extended Data Figure 3.**
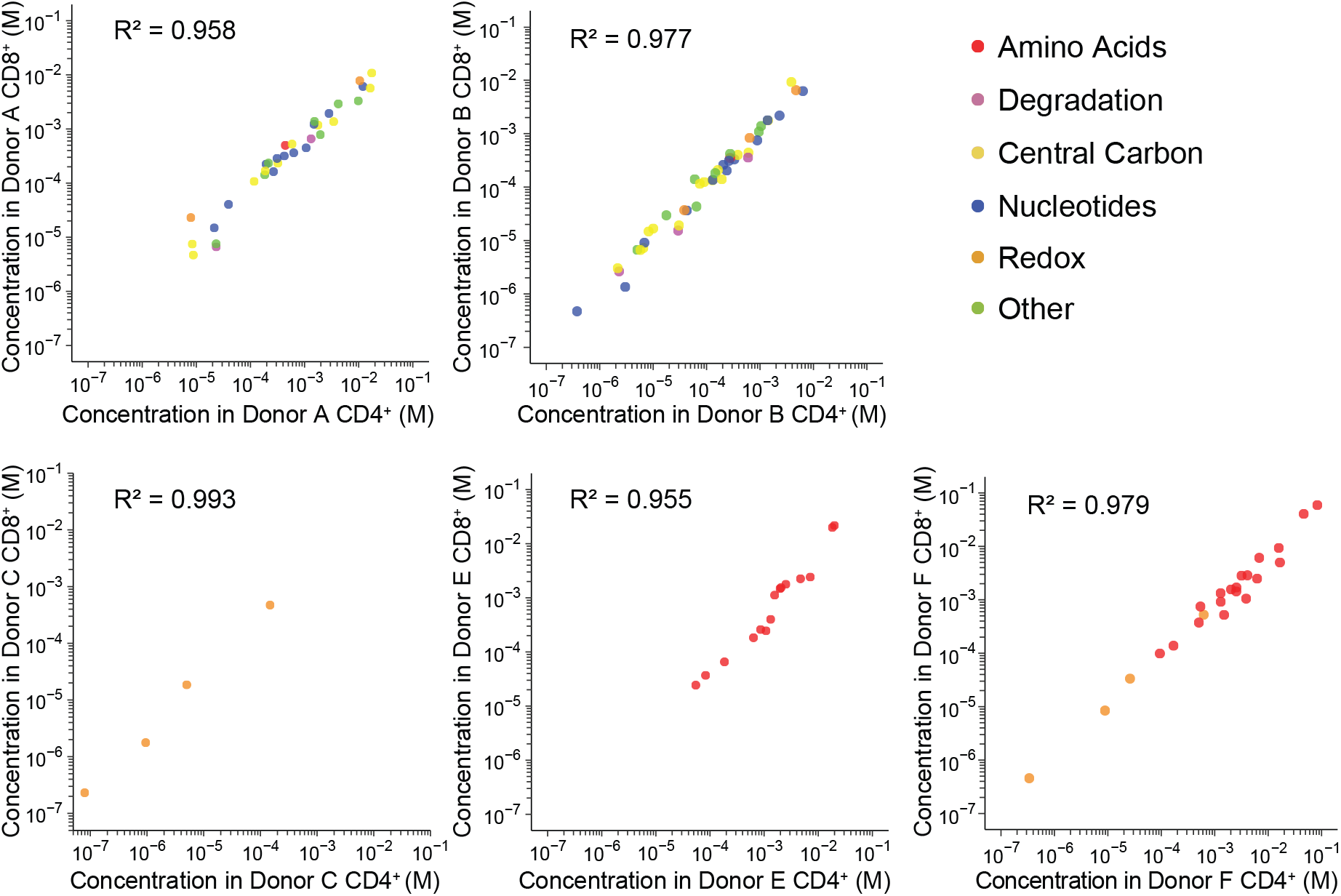
Comparison of T cell metabolomes across subtypes from the same individuals. Absolute metabolite concentrations in CD4^+^ helper T cells and CD8^+^ killer T cells from the same individuals were compared.

**Extended Data Figure 4.**
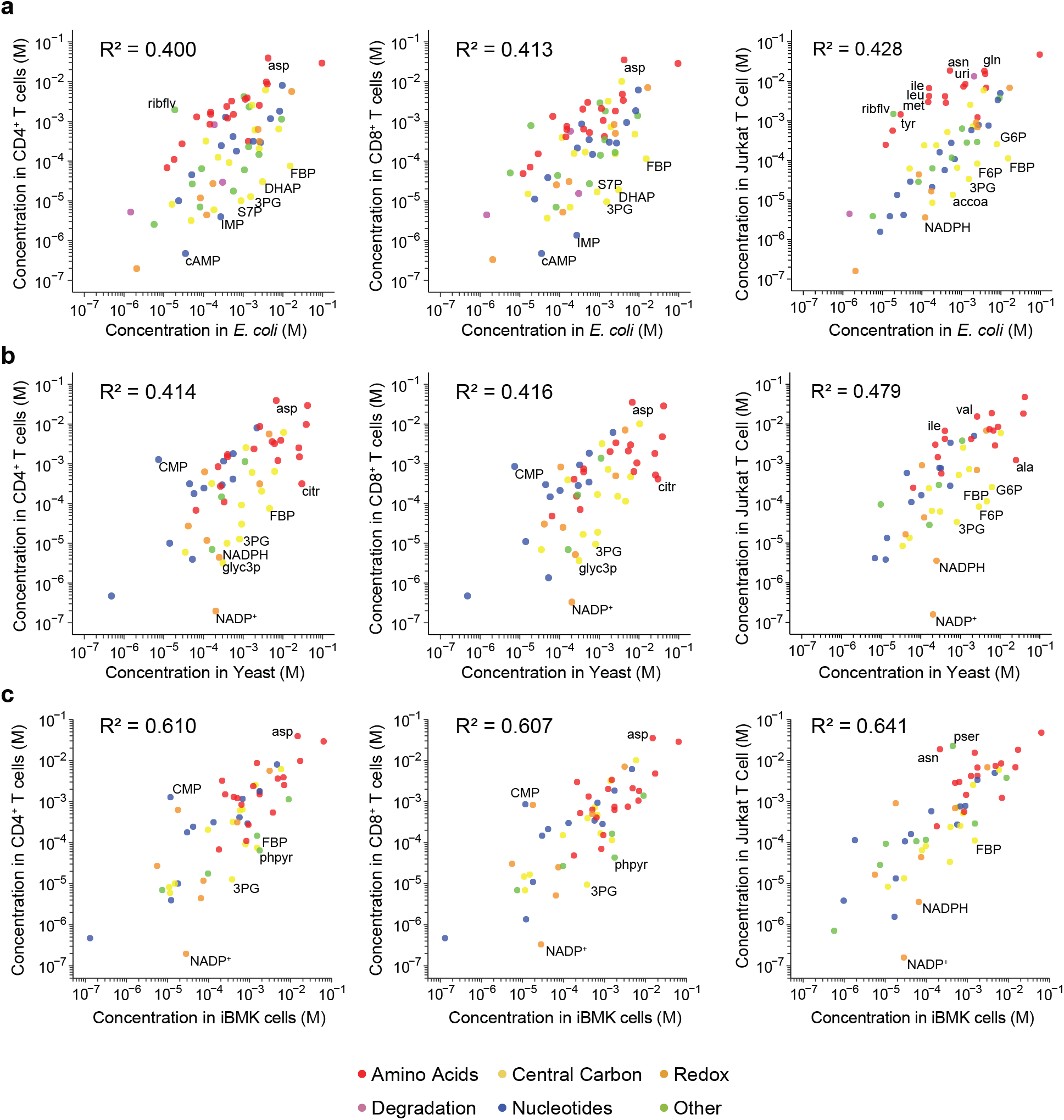
Comparison of metabolomes between divergent cell types. (**a-c**) Absolute metabolite concentrations in CD4^+^, CD8^+^ and Jurkat T cells are compared to those of (**a**) *E. coli*, (**b**) yeast, and (**c**) iBMK cells. G6P denotes glucose-6-phosphate; F6P, fructose-6-phosphate; FBP, fructose-1,6-bisphosphate; 3PG, 3-phosphoglycerate; glyc3p, glycerol-3-phosphate; phpyr, phenylpyruvate; citr, citrulline; pser, phosphoserine; ribflv, riboflavin; uri, uridine; and accoa, acetyl-CoA.

**Extended Data Figure 5.**
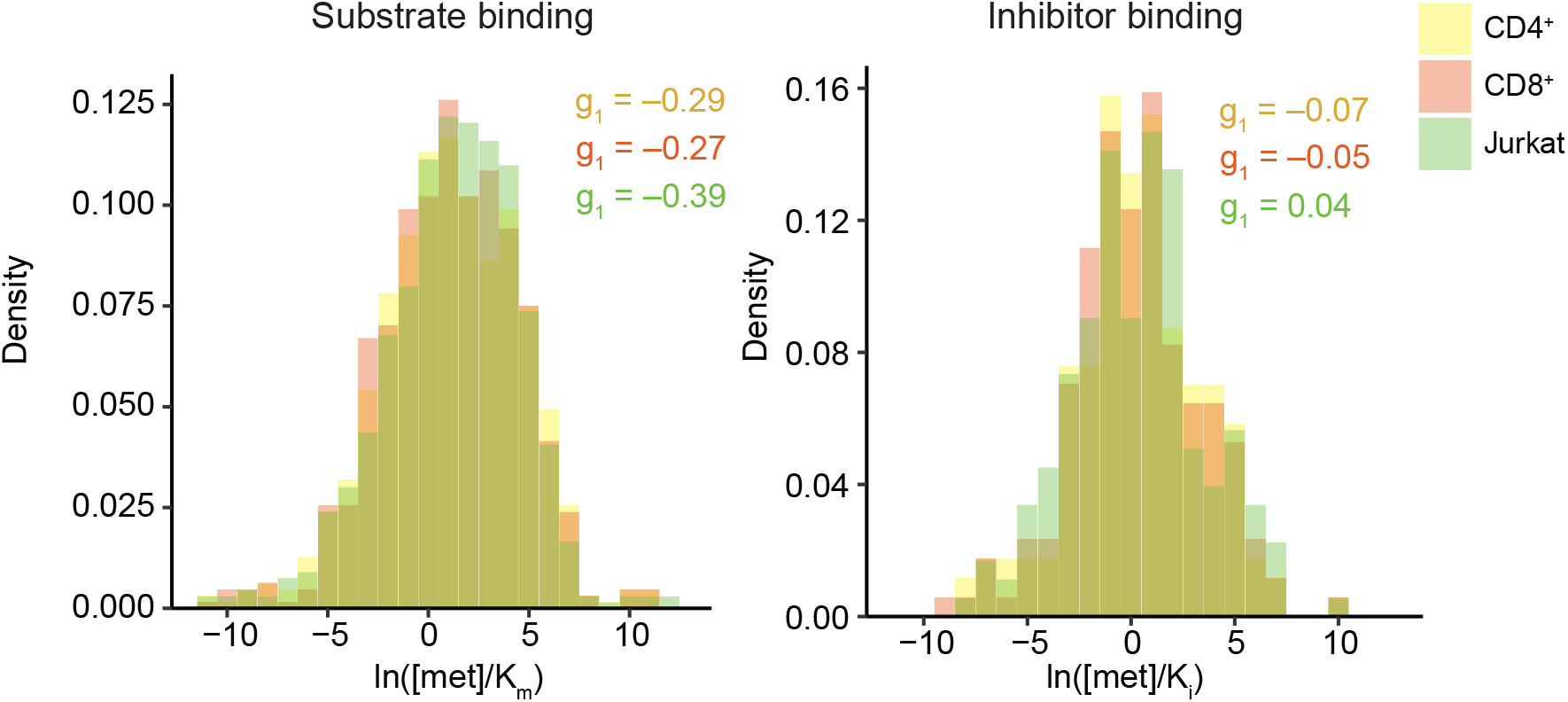
Conservation of substrate and inhibitor binding site occupancy across T cell types. The Kolmogorov-Smirnov test on log ratios of metabolite concentrations to K_m_ and K_i_ concluded no significant differences between T cell types. g_1_ represents the Fisher-Pearson moment coefficient of skewness for CD4^+^ (yellow), CD8^+^ (red), and Jurkat (green) T cells.

**Extended Data Figure 6.**
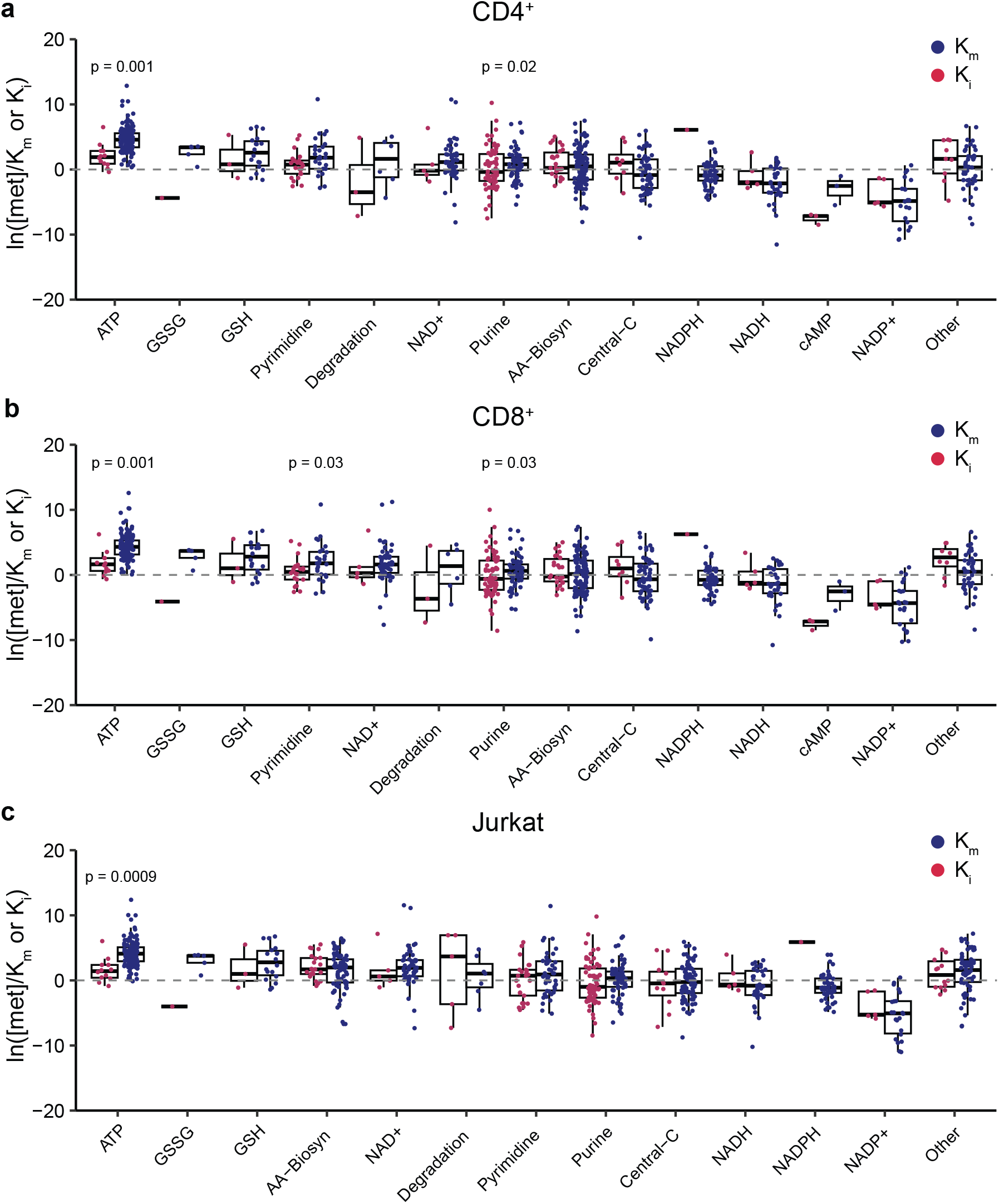
Comparison of substrate and inhibitor binding site occupancy across metabolite groups. (**a-c**) Substrate and inhibitor binding site occupancies were compared for individual metabolite groups in (**a**) CD4^+^, (**b**) CD8^+^, and (**c**) Jurkat T cells. Boxplots show the quartiles with thick lines representing the median. The whiskers extend to the farthest measurement values within 1.5 times the IQR from the edges of the box. Statistical significance between substrate and inhibitor binding site occupancies for each metabolite group was assessed by the Kolmogorov-Smirnov test and multiple comparison correction using the Benjamini-Hochberg procedure.

**Extended Data Figure 7.**
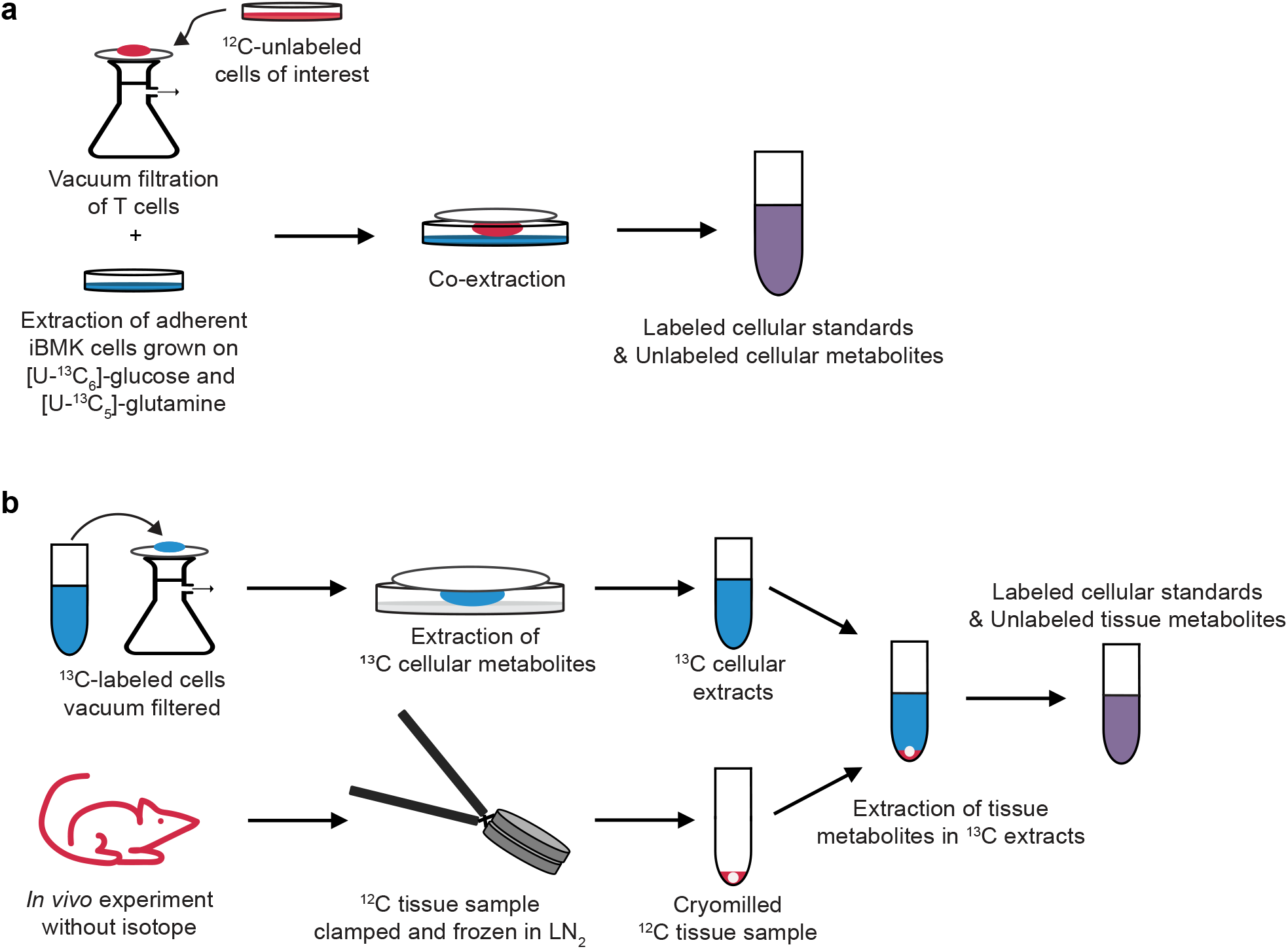
Metabolite co-extraction methods for adherent cells and tissues. (**a**) Adherent cells (e.g., iBMK cells) are extracted in the plate that they are cultured, and suspension cells (e.g., T cells) are extracted in the same plate after vacuum filtration. (**b**) Tissues are clamped, snap frozen in liquid nitrogen (LN_2_), and cryomilled before metabolite extraction using the extracts of ^13^C-labeled reference cells.

**Extended Data Figure 8.**
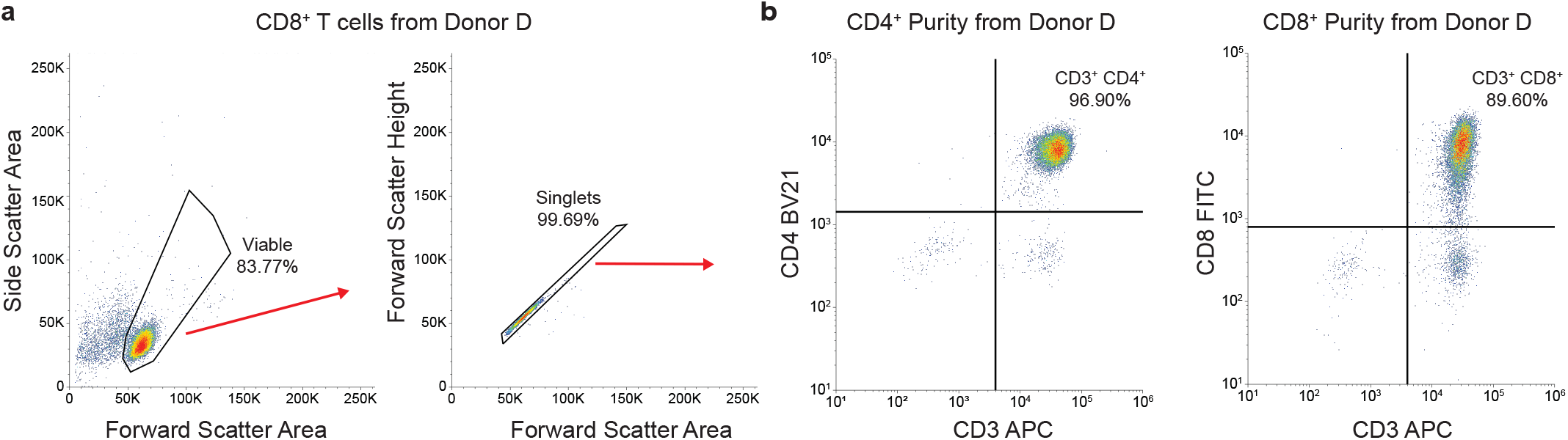
Gating strategy for selecting viable cells and determining their subtypes. (**a**) Initial gating excluded debris by analyzing forward-scatter area (FSC-A) and side-scatter area (SSC-A) parameters. Doublets were then excluded using forward-scatter area (FSC-A) versus forward-scatter height (FSC-H). (**b**) The purity of the cell population was assessed by evaluating the singlets based on the fluorescence intensity of anti-CD4 and anti-CD8 antibody staining in conjunction with anti-CD3 antibody staining.

**Extended Data Figure 9.**
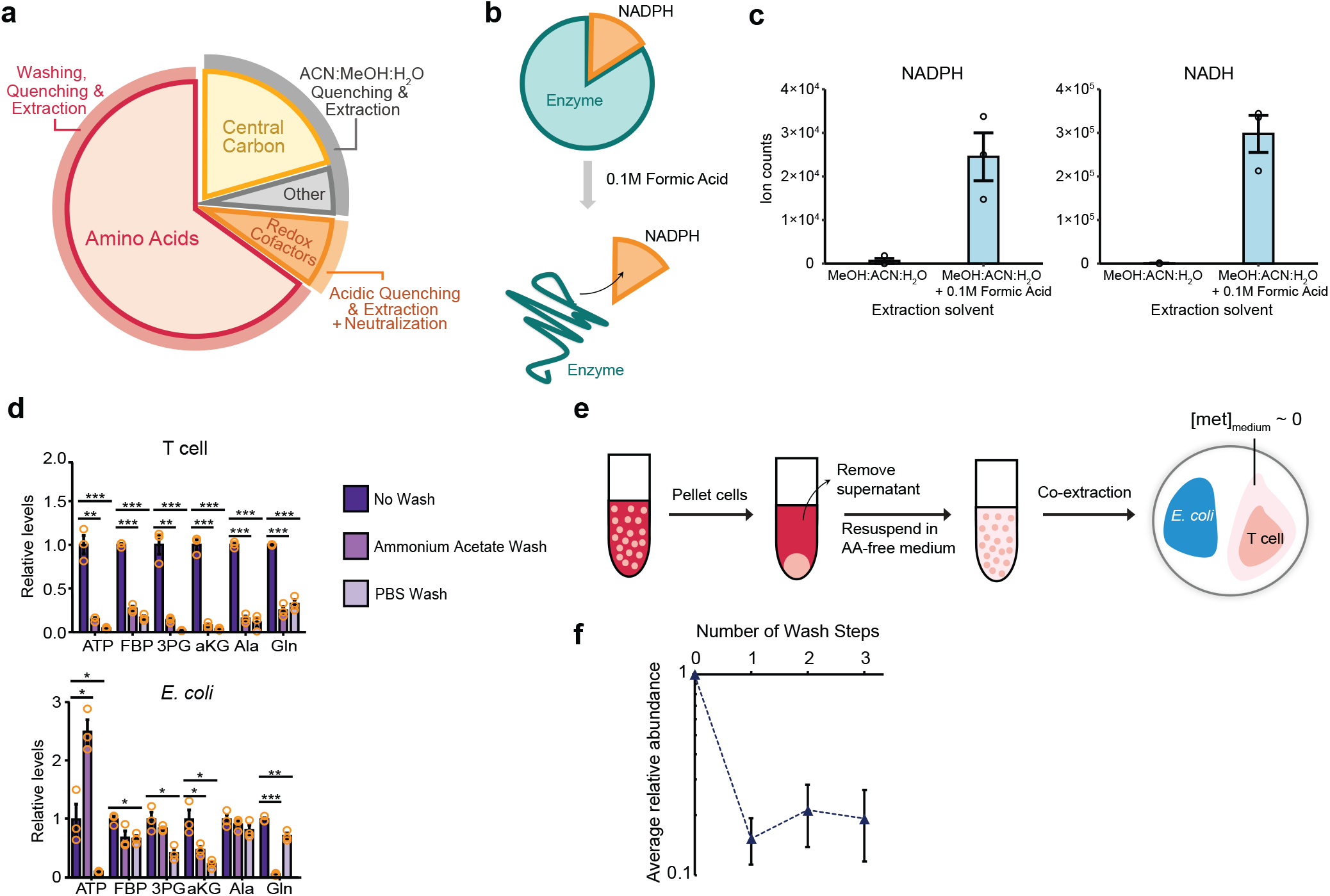
Tailored extraction methods capture a broad range of metabolites. (**a**) Three methods for quenching metabolism and extracting metabolite, which were tailored to different metabolite classes, enabled comprehensive metabolite quantitation. (**b**) To extract and measure NAD(P)H, an acidic extraction solvent containing 0.1M formic acid was used to denature enzymes and dissociate them. Acidic extracts were neutralized to prevent the degradation of metabolites. (**c**) Acidic quenching increased ion counts for NADPH and NADH. Error bars represent the s.e.m. (n = 3 biological replicates). (**d**) Relative abundances of representative metabolites were measured with and without washing cells directly on filters. Either 150 mM ammonium acetate (pH 7.4) or PBS was used for washing T cells and *E. coli*. Error bars represent the s.e.m. (n = 3 replicates). P values were calculated using two-tailed t-tests (* represents p<0.05, ** represents p<0.01, and *** represents p<0.001). (**e**) To measure intracellular amino acids without interference from extracellular amino acids, cells were pelleted, supernatant was quickly removed, and cells were resuspended in an amino acid-free medium prior to co-extraction. (**f**) Amino acids were measured after wash steps and compared to the measurement with no wash. The averages of relative amino acid levels are shown with error bars representing the s.e.m. (n = 14 amino acids). A single centrifuge-and-wash step was sufficient to eliminate the interference from extracellular amino acids.

## References

1 Biasco, L. et al. Clonal expansion of T memory stem cells determines early anti-leukemic responses and long-term CAR T cell persistence in patients. Nature cancer 2, 629–642 (2021).

2 Liu, C., Qi, T., Milner, J. J., Lu, Y. & Cao, Y. Speed and location both matter: antigen stimulus dynamics controls CAR-T cell response. Frontiers in Immunology 12, 748768 (2021).

3 Acha-Sagredo, A. et al. A constitutive interferon-high immunophenotype defines response to immunotherapy in colorectal cancer. Cancer Cell 43, 292-307. e297 (2025).

4 Walsh, Z. H. et al. Mapping variant effects on anti-tumor hallmarks of primary human T cells with base-editing screens. Nature biotechnology 43, 384–395 (2025).

5 Quast, N. P. et al. T-cell receptor structures and predictive models reveal comparable alpha and beta chain structural diversity despite differing genetic complexity. Communications Biology 8, 362 (2025).

6 Chang, J. T. et al. Asymmetric T lymphocyte division in the initiation of adaptive immune responses. science 315, 1687–1691 (2007).

7 Macintyre, A. N. et al. The glucose transporter Glut1 is selectively essential for CD4 T cell activation and effector function. Cell metabolism 20, 61–72 (2014).

8 Zeng, H. et al. mTORC1 and mTORC2 kinase signaling and glucose metabolism drive follicular helper T cell differentiation. Immunity 45, 540–554 (2016).

9 Michalek, R. D. et al. Cutting edge: distinct glycolytic and lipid oxidative metabolic programs are essential for effector and regulatory CD4+ T cell subsets. The Journal of immunology 186, 3299–3303 (2011).

10 Klein Geltink, R. I. et al. Metabolic conditioning of CD8(+) effector T cells for adoptive cell therapy. Nat Metab 2, 703–716 (2020).

11 Klein Geltink, R. I., Kyle, R. L. & Pearce, E. L. Unraveling the complex interplay between T cell metabolism and function. Annual review of immunology 36, 461–488 (2018).

12 Mendoza, A. et al. Lymphatic endothelial S1P promotes mitochondrial function and survival in naive T cells. Nature 546, 158–161 (2017).

13 Shi, Y., Zhang, H. & Miao, C. Metabolic reprogram and T cell differentiation in inflammation: current evidence and future perspectives. Cell Death Discovery 11, 123 (2025).

14 Lakhani, A. et al. Extracellular domains of CARs reprogramme T cell metabolism without antigen stimulation. Nature Metabolism, 1–18 (2024).

15 Wang, R. et al. The transcription factor Myc controls metabolic reprogramming upon T lymphocyte activation. Immunity 35, 871–882 (2011).

16 Sena, L. A. et al. Mitochondria are required for antigen-specific T cell activation through reactive oxygen species signaling. Immunity 38, 225–236 (2013).

17 Laniewski, N. G. & Grayson, J. M. Antioxidant treatment reduces expansion and contraction of antigen-specific CD8+ T cells during primary but not secondary viral infection. Journal of virology 78, 11246–11257 (2004).

18 Morris, G., Gevezova, M., Sarafian, V. & Maes, M. Redox regulation of the immune response. Cellular & Molecular Immunology 19, 1079–1101 (2022).

19 Tavassolifar, M. j., Vodjgani, M., Salehi, Z. & Izad, M. The influence of reactive oxygen species in the immune system and pathogenesis of multiple sclerosis. Autoimmune diseases 2020, 5793817 (2020).

20 Metallo, C. M. & Vander Heiden, M. G. Understanding metabolic regulation and its influence on cell physiology. Molecular cell 49, 388–398 (2013).

21 Park, J. O. et al. Near-equilibrium glycolysis supports metabolic homeostasis and energy yield. Nature chemical biology 15, 1001–1008 (2019).

22 Wegner, A., Meiser, J., Weindl, D. & Hiller, K. How metabolites modulate metabolic flux. Current opinion in biotechnology 34, 16–22 (2015).

23 Bennett, B. D. et al. Absolute metabolite concentrations and implied enzyme active site occupancy in Escherichia coli. Nature chemical biology 5, 593–599 (2009).

24 Park, J. O. et al. Metabolite concentrations, fluxes and free energies imply efficient enzyme usage. Nature chemical biology 12, 482–489 (2016).

25 Alseekh, S. et al. Mass spectrometry-based metabolomics: a guide for annotation, quantification and best reporting practices. Nature methods 18, 747–756 (2021).

26 Guder, J. C., Schramm, T., Sander, T. & Link, H. Time-optimized isotope ratio LC–MS/MS for high-throughput quantification of primary metabolites. Analytical Chemistry 89, 1624–1631 (2017).

27 Bennett, B. D., Yuan, J., Kimball, E. H. & Rabinowitz, J. D. Absolute quantitation of intracellular metabolite concentrations by an isotope ratio-based approach. Nature protocols 3, 1299–1311 (2008).

28 Ma, E. H. et al. Metabolic profiling using stable isotope tracing reveals distinct patterns of glucose utilization by physiologically activated CD8+ T cells. Immunity 51, 856–870. e855 (2019).

29 Scherer, S. et al. Pyrimidine de novo synthesis inhibition selectively blocks effector but not memory T cell development. Nature immunology 24, 501–515 (2023).

30 Mora, T., Walczak, A. M., Bialek, W. & Callan Jr, C. G. Maximum entropy models for antibody diversity. Proceedings of the National Academy of Sciences 107, 5405–5410 (2010).

31 Brunk, E. et al. Recon3D enables a three-dimensional view of gene variation in human metabolism. Nature biotechnology 36, 272–281 (2018).

32 Wu, B. et al. Mitochondrial aspartate regulates TNF biogenesis and autoimmune tissue inflammation. Nature immunology 22, 1551–1562 (2021).

33 Zhao, H. et al. Myeloid-derived itaconate suppresses cytotoxic CD8+ T cells and promotes tumour growth. Nature metabolism 4, 1660–1673 (2022).

34 Wang, H. et al. Aspartate metabolism facilitates IL-1β production in inflammatory macrophages. Frontiers in immunology 12, 753092 (2021).

35 Jha, A. K. et al. Network integration of parallel metabolic and transcriptional data reveals metabolic modules that regulate macrophage polarization. Immunity 42, 419–430 (2015).

36 Teh, M. R. et al. Iron deficiency causes aspartate-sensitive dysfunction in CD8+ T cells. Nature Communications 16, 5355 (2025).

37 Xia, W., Mao, Y., Xia, Z., Cheng, J. & Jiang, P. Metabolic remodelling produces fumarate via the aspartate– argininosuccinate shunt in macrophages as an antiviral defence. Nature Microbiology 10, 1115–1129 (2025).

38 Noor, E. et al. Pathway thermodynamics highlights kinetic obstacles in central metabolism. PLoS computational biology 10, e1003483 (2014).

39 Chapman, A. G., Fall, L. & Atkinson, D. E. Adenylate energy charge in Escherichia coli during growth and starvation. Journal of bacteriology 108, 1072–1086 (1971).

40 Mak, T. W. et al. Glutathione primes T cell metabolism for inflammation. Immunity 46, 675–689 (2017).

41 Romeu, M. et al. Evaluation of oxidative stress biomarkers in patients with chronic renal failure: a case control study. BMC research notes 3, 1–7 (2010).

42 Giustarini, D. et al. Pitfalls in the analysis of the physiological antioxidant glutathione (GSH) and its disulfide (GSSG) in biological samples: An elephant in the room. Journal of Chromatography B 1019, 21–28 (2016).

43 Enns, G. M. et al. Degree of glutathione deficiency and redox imbalance depend on subtype of mitochondrial disease and clinical status. PloS one 9, e100001 (2014).

44 Valsecchi, F. et al. Metabolic consequences of NDUFS4 gene deletion in immortalized mouse embryonic fibroblasts. Biochimica et Biophysica Acta (BBA)-Bioenergetics 1817, 1925–1936 (2012).

45 Buck, M. D., O’sullivan, D. & Pearce, E. L. T cell metabolism drives immunity. Journal of Experimental Medicine 212, 1345–1360 (2015).

46 Buck, M. D. et al. Mitochondrial dynamics controls T cell fate through metabolic programming. Cell 166, 63–76 (2016).

47 Chang, C.-H. et al. Posttranscriptional control of T cell effector function by aerobic glycolysis. Cell 153, 1239–1251 (2013).

48 Greiner, E. F., Guppy, M. & Brand, K. Glucose is essential for proliferation and the glycolytic enzyme induction that provokes a transition to glycolytic energy production. Journal of Biological Chemistry 269, 31484–31490 (1994).

49 Pearce, E. L., Poffenberger, M. C., Chang, C.-H. & Jones, R. G. Fueling immunity: insights into metabolism and lymphocyte function. Science 342, 1242454 (2013).

50 Reina-Campos, M., Scharping, N. E. & Goldrath, A. W. CD8+ T cell metabolism in infection and cancer. Nature Reviews Immunology 21, 718–738 (2021).

51 Kishton, R. J., Sukumar, M. & Restifo, N. P. Metabolic regulation of T cell longevity and function in tumor immunotherapy. Cell metabolism 26, 94–109 (2017).

52 Siska, P. J. et al. Suppression of Glut1 and glucose metabolism by decreased Akt/mTORC1 signaling drives T cell impairment in B cell leukemia. The Journal of Immunology 197, 2532–2540 (2016).

53 Crompton, J. G. et al. Akt inhibition enhances expansion of potent tumor-specific lymphocytes with memory cell characteristics. Cancer research 75, 296–305 (2015).

54 Franchi, L. et al. Inhibiting oxidative phosphorylation in vivo restrains Th17 effector responses and ameliorates murine colitis. The Journal of Immunology 198, 2735–2746 (2017).

55 Zhao, Q. et al. CD4+ T cell activation and concomitant mTOR metabolic inhibition can ablate microbiota-specific memory cells and prevent colitis. Science immunology 5, eabc6373 (2020).

56 Yin, Y. et al. (View at Publisher| View at, 2015).

57 Hackett, S. R. et al. Systems-level analysis of mechanisms regulating yeast metabolic flux. Science 354, aaf2786 (2016).

58 Fell, D. & Cornish-Bowden, A. Understanding the control of metabolism. Vol. 2 (Portland press London, 1997).

59 Hope, H. C. et al. Coordination of asparagine uptake and asparagine synthetase expression modulates CD8+ T cell activation. JCI insight 6, e137761 (2021).

60 Wu, J. et al. Asparagine enhances LCK signalling to potentiate CD8+ T-cell activation and anti-tumour responses. Nature cell biology 23, 75–86 (2021).

61 Gnanaprakasam, J. R. et al. Asparagine restriction enhances CD8+ T cell metabolic fitness and antitumoral functionality through an NRF2-dependent stress response. Nature Metabolism 5, 1423–1439 (2023).

62 Mills, E. L. et al. Succinate dehydrogenase supports metabolic repurposing of mitochondria to drive inflammatory macrophages. Cell 167, 457–470. e413 (2016).

63 Mills, E. L. et al. Itaconate is an anti-inflammatory metabolite that activates Nrf2 via alkylation of KEAP1. Nature 556, 113–117 (2018).

64 Seim, G. L. et al. Two-stage metabolic remodelling in macrophages in response to lipopolysaccharide and interferon-γ stimulation. Nature metabolism 1, 731–742 (2019).

65 Young, J. D. INCA: a computational platform for isotopically non-stationary metabolic flux analysis. Bioinformatics 30, 1333–1335 (2014).

66 O’Keeffe, S. et al. Bringing carbon to life via one-carbon metabolism. Trends in Biotechnology (2024).

67 Chen, W. W., Freinkman, E., Wang, T., Birsoy, K. & Sabatini, D. M. Absolute quantification of matrix metabolites reveals the dynamics of mitochondrial metabolism. Cell 166, 1324–1337. e1311 (2016).

68 Piazza, I. et al. A map of protein-metabolite interactions reveals principles of chemical communication. Cell 172, 358–372. e323 (2018).

69 Dixit, A. & Kalia, J. Protein-Metabolite Interactions: Discovery and Significance. ChemBioChem 24, e202200755 (2023).

70 Kelley, L. P. et al. Integrated analysis of transcriptional and metabolic responses to mitochondrial stress. Cell Reports Methods 5 (2025).

71 Hatton, I. A. et al. The human cell count and size distribution. Proceedings of the National Academy of Sciences 120, e2303077120 (2023).

72 Stanciu, L. A., Shute, J., Holgate, S. T. & Djukanovic, R. Production of IL-8 and IL-4 by positively and negatively selected CD4+ and CD8+ human T cells following a four-step cell separation method including magnetic cell sorting (MACS). Journal of immunological methods 189, 107–115 (1996).

73 Gutnick, D., Calvo, J. M., Klopotowski, T. & Ames, B. N. Compounds which serve as the sole source of carbon or nitrogen for Salmonella typhimurium LT-2. Journal of bacteriology 100, 215–219 (1969).

74 Degenhardt, K. & White, E. A mouse model system to genetically dissect the molecular mechanisms regulating tumorigenesis. Clinical cancer research 12, 5296–5304 (2006).

75 Rabinowitz, J. D. & Kimball, E. Acidic acetonitrile for cellular metabolome extraction from Escherichia coli. Analytical chemistry 79, 6167–6173 (2007).

76 Wang, L. et al. Peak annotation and verification engine for untargeted LC–MS metabolomics. Analytical chemistry 91, 1838–1846 (2018).

77 Seitzer, P., Bennett, B. & Melamud, E. MAVEN2: an updated open-source mass spectrometry exploration platform. Metabolites 12, 684 (2022).

78 Schomburg, I. et al. BRENDA in 2013: integrated reactions, kinetic data, enzyme function data, improved disease classification: new options and contents in BRENDA. Nucleic acids research 41, D764–D772 (2012).

79 Flamholz, A., Noor, E., Bar-Even, A. & Milo, R. eQuilibrator—the biochemical thermodynamics calculator. Nucleic acids research 40, D770–D775 (2012).

80 Xu, H., Yang, D., Jiang, D. & Chen, H.-Y. Phosphate assay kit in one cell for electrochemical detection of intracellular phosphate ions at single cells. Frontiers in Chemistry 7, 360 (2019).

81 Bevington, A. et al. A study of intracellular orthophosphate concentration in human muscle and erythrocytes by 31P nuclear magnetic resonance spectroscopy and selective chemical assay. Clinical Science 71, 729–735 (1986).

82 Behjousiar, A., Kontoravdi, C. & Polizzi, K. M. In situ monitoring of intracellular glucose and glutamine in CHO cell culture. PLoS One 7, e34512 (2012).

83 Bearham, J., Garnett, J. P., Schroeder, V., Biggart, M. G. & Baines, D. L. Effective glucose metabolism maintains low intracellular glucose in airway epithelial cells after exposure to hyperglycemia. American Journal of Physiology-Cell Physiology 317, C983–C992 (2019).

84 Xu, J., Huang, P., Qin, Y., Jiang, D. & Chen, H.-y. Analysis of intracellular glucose at single cells using electrochemiluminescence imaging. Analytical chemistry 88, 4609–4612 (2016).

